# Mapping the genetic landscape of cell morphogenesis and subcellular organization in stationary-phase *Escherichia coli* cells

**DOI:** 10.64898/2026.11.13.766414

**Authors:** Bart Steemans, Kaat Sondervorst, Matthew Herdman, Kristina Nesporova, Alix Meunier, Els Meert, Sander K. Govers

## Abstract

In nature, many bacteria reside in a non-growing stationary phase. The extent and genetic basis of the accompanying cellular remodeling remain unclear. Here, we developed Coli-Painting, a combinatorial staining assay, and BactoScoop, a dedicated image-based profiling pipeline, to characterize the growth-to-stasis transition in *Escherichia coli*. Single-cell profiles revealed a gradual yet heterogeneous entry, with some cells retaining signs of chromosome replication or cell division well into stationary phase. Subsequent genome-wide profiling identified genetic determinants of stationary-phase cell morphology, nucleoid organization, RNA distribution, and membrane-related properties. Global correlations among cellular features revealed coordinated aspects of stationary-phase cell architecture, while high-dimensional mutant profiles provided functional context for genes of known and unknown function. Downstream characterization of 19 y-genes highlighted our approach’s ability to identify relevant contributors to stationary-phase physiology. Overall, our work maps the genetic landscape of stationary-phase cell organization in *E. coli* and serves as a resource for uncovering growth-stage-specific gene function.

## Main

Bacterial populations frequently experience nutrient limitation and environmental stress in nature, prompting a transition from active growth to a quiescent stationary phase^1,2^. This transition represents a highly regulated physiological reprogramming that promotes long-term survival and broad stress resistance, including tolerance to antibiotics^3–5^. Extensive studies in the model bacterium *Escherichia coli* have documented the biochemical hallmarks of this shift, including a global slowdown of growth and protein synthesis, metabolic reorientation toward maintenance functions, and activation of broad stress-response programs^6–16^. This process is largely driven by the activation of specific regulatory programs such as the general stress response (mediated by the general stress sigma factor RpoS^17,18^) and the stringent response (mediated by alarmone (p)ppGpp levels^19,20^). While the biochemical remodeling of stationary-phase cells is increasingly well defined, their accompanying structural transformation remains less understood. In *E. coli*, average cell size decreases upon entry into stationary phase through reductive cell divisions, in which initiated rounds of chromosome replication and subsequent cell division are completed in the absence of cell growth^5,21^. This process leads to an increase in the surface-area-to-volume ratio of cells^22,23^ that is believed to maximize nutrient acquisition and waste product excretion. In addition, nucleoids are organized into dense, ordered structures offering protection against oxidative and mutagenic damage, a process that is mediated, at least in part, by nucleoid-associated proteins such as Dps and H-NS^5,24–28^. The biophysical properties of the cytoplasm are also altered in stationary phase^29^, the structural integrity of the cell envelope changes^30,31^, and the periplasm increases in size (at the expense of the cytoplasm)^8,32^. Together, these observations suggest a broad cellular reorganization, yet the exact changes and their underlying genetic determinants are mostly unresolved.

Adding further complexity to this structural transformation is the fact that entry into stationary phase does not occur uniformly^21,33^, nor does stationary phase itself represent a homogeneous and time-invariant assembly of bacterial cells. Instead, isogenic stationary populations consist of a dynamic and heterogeneous mixture of cells, displaying striking heterogeneity in size^21^, nucleoid compaction^34^, cell division^35^, and potential for growth resumption^21,33,36^. Such heterogeneity had previously remained undetected in bulk-level population assays^37,38^, but has been uncovered in more recent years through various single-cell approaches. While this phenotypic heterogeneity likely reflects an evolutionary bet-hedging strategy to generate subpopulations optimized for surviving distinct environments (e.g., increased tolerance to antibiotics^21,39^ or prolonged survival during starvation^40,41^), we do not understand its exact extent nor how it arises as populations transition to stationary phase.

Here, we address these unknowns by providing an in-depth characterization of the cellular reorganization that occurs in *E. coli* upon entering stationary phase at an unprecedented scale and resolution. To achieve this, we developed a combinatorial staining assay to visualize different cellular components (named Coli-Painting, in analogy to the Cell Painting assays that exist for eukaryotic cells^42^), a multispectral imaging protocol, and a dedicated image-based profiling pipeline (named BactoScoop). The BactoScoop pipeline combines image analysis and machine learning methods to provide high-content cell biological information in the form of >650 single-cell level features related to cell morphology and cellular organization. Using this approach, we carried out a high-throughput image-based screen of the *E. coli* single-gene knockout collection during stationary phase, imaging 3,703 mutants and generating a dataset of >15.8 million individual cells. This comprehensive analysis enabled us to determine single-cell level heterogeneity upon entering and within stationary phase, and identified many genes previously unrecognized for their roles in stationary-phase cell morphogenesis and subcellular organization, including several genes of unknown function.

## Results

### Cell biological features related to cell morphology, cell division, nucleoid morphology, and chromosome replication plateau in *E. coli* upon entry into stationary phase

Given the dynamic and heterogeneous nature of stationary-phase populations, we first set out to characterize single-cell behavior along the growth curve of *E. coli* populations. For this, we used an *E. coli* strain carrying a chromosomally encoded GFP fusion to DnaN (i.e., the β sliding clamp of DNA polymerase III) to monitor DNA replication^43^. This strain was grown in M9 medium supplemented with 0.2% glucose (M9glu) and live cells were sampled for phase-contrast and fluorescence microscopy from exponential phase to late stationary phase (Fig. 1a-b). Nucleoids were visualized by staining cells with the DNA dye 4’,6-diamidino-2-phenylindole (DAPI). At each sampling point, we extracted single-cell features related to cell morphology, cell division, nucleoid morphology and chromosome replication (Dataset S1), and we determined the fraction of cells displaying cell constriction, containing a single nucleoid, and performing chromosome replication (Extended Data Fig. 1a-b, Methods).

**Fig. 1:**
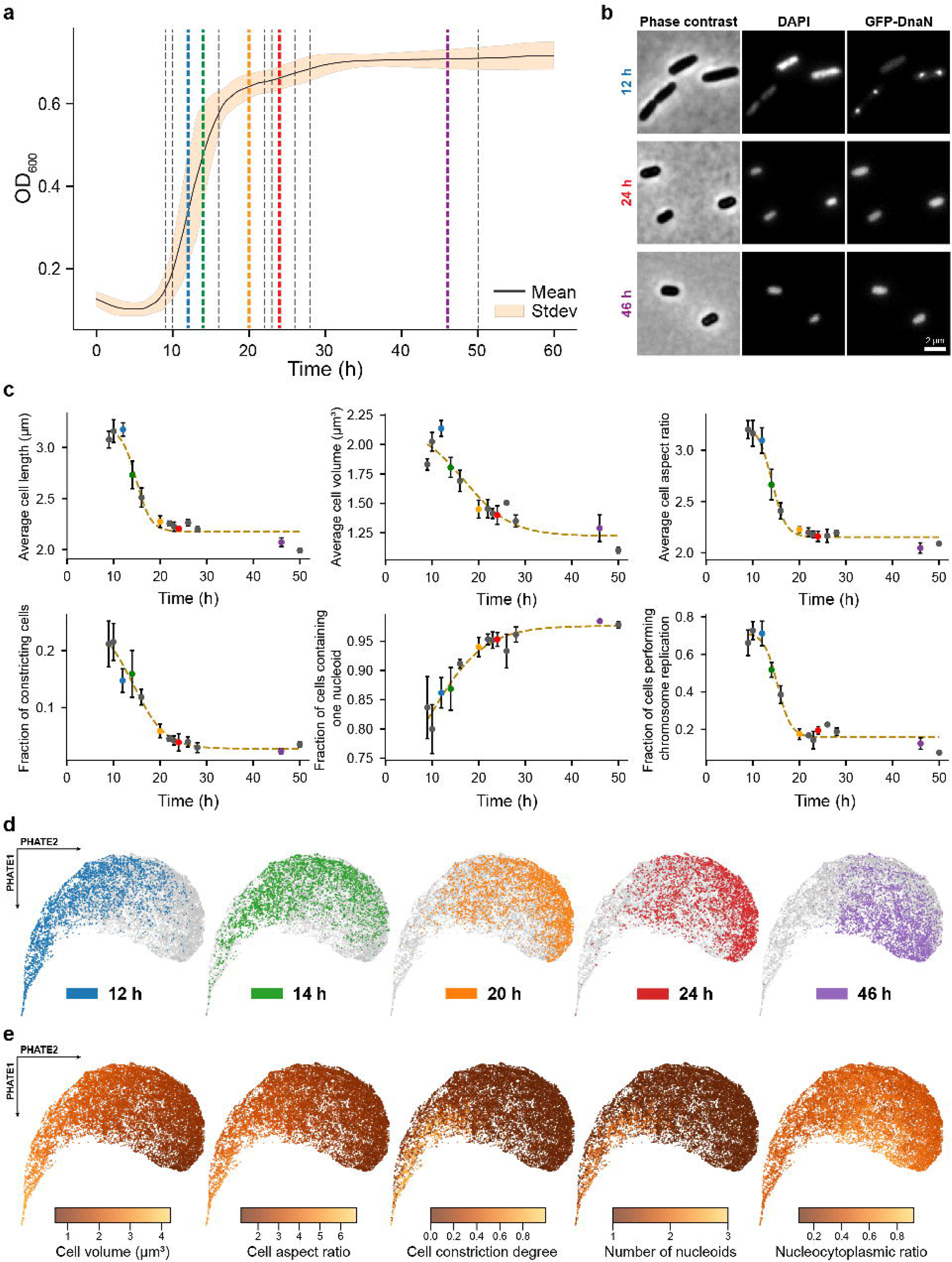
Cell morphological and replicative characteristics reach a plateau upon entry into stationary phase. **a**, Growth curve of *E. coli* OZA001^43^ in M9 medium supplemented with 0.2% glucose. Dashed lines indicate sampling times of the time-course experiment. **b,** Representative phase-contrast and epifluorescence images of cells at timepoints 12 h, 24 h, and 46 h. **c,** Plots showing the behavior of average cell morphological features (cell length, volume, and aspect ratio) and cell cycle features (fraction of constricting cells, cells with one nucleoid, and fraction of cells performing chromosome replication) over time. Error bars indicate standard deviation between replicates (n ≥ 3). **d,** Feature embedding of 13 cell and nucleoid morphological features across five timepoints. Cells from each timepoint are highlighted separately in the indicated color. **e,** The feature embedding of **d** overlaid with cell morphological and nucleoid features.

Consistent with previous studies^22,44,45^, average cellular properties changed markedly along the growth curve. Growth-related cell morphological features, such as average cell length and volume, decreased progressively during nutrient depletion and entry into stationary phase, seemingly stabilizing as cells reached deeper stages of stationary phase (Fig. 1c). We observed a similar trend for the fraction of constricting cells and the fraction of cells engaged in chromosome replication (Fig. 1c). At the same time, the fraction of cells with a single nucleoid gradually increased and stabilized deeper into stationary phase (Fig. 1c). Cell width only displayed minor changes and remained largely constant (Extended Data Fig. 1c), leading to the cell aspect ratio displaying a similar pattern as cell length and volume (Fig. 1c). While total cell surface area displayed a similar pattern to cell length and volume (Extended Data Fig. 1c), the surface-area-to-volume ratio increased upon entering stationary phase (Extended Data Fig. 1c), in line with previous observations^22,23^, indicating that the decrease in cell surface area is less pronounced than that of cell volume. These observations suggest that cells gradually arrest their growth upon entry into stationary phase and maintain that growth arrest within stationary phase. However, while the fraction of cells displaying growth-related properties decreased, it never reached zero. At all sampling points, at least 1.54% of the population retained detectable signs of cell division and/or chromosome replication and segregation, even deep into stationary phase (50 h after reinoculation; Fig. 1c). We made similar observations in an undefined nutrient-rich growth medium (LB), although the timing and density at which entry into stationary phase occurred differed (Extended Data Fig. 1d-e, Dataset S1). Together, these findings indicate an asynchronous and incomplete growth arrest of the population upon entry into stationary phase, across different initial growth conditions.

In order to visualize single-cell heterogeneity across the growth curve, we applied PHATE (Potential of Heat-diffusion for Affinity-based Transition Embedding^46^) to a set of 13 cell- and nucleoid-morphology features extracted from individual cells at five timepoints (Dataset S1). The resulting embedding (combining cells from each timepoint) captures the continuum of morphological diversity both within and between these populations and forms a gradual trajectory that reflects the smooth yet heterogeneous morphological transition from exponential growth to stationary phase, with substantial population overlap between timepoints (Fig. 1d). Projecting individual features onto the embedding further clarifies this trajectory and highlights how their behavior changes towards and within stationary phase (Fig. 1e). These findings align with our previous observations of average cellular features (Fig. 1c) and further underscore the gradual entry into stationary phase while also highlighting the heterogeneous nature of the growth-to-stasis transition at the single-cell level.

### Coli-Painting and BactoScoop image-based profiling enable the high-throughput phenotyping of stationary-phase *E. coli* cells

To better assess the structural changes of stationary-phase *E. coli* cells and heterogeneity therein, we developed the Coli-Painting assay and BactoScoop image-based profiling pipeline. The Coli-Painting assay uses spectrally compatible and commercially available dyes to simultaneously visualize several major cellular components (i.e., RNA, DNA, inner membrane, outer membrane), alongside cell morphology (Fig. 2a). Our tailored BactoScoop image-based profiling pipeline subsequently extracts quantitative single-cell level features from the obtained microscopy images (Fig. 2b, Methods). Combining both approaches enables automated high-throughput data collection across thousands of cells, each described by >650 cell biological features (i.e., a profile).

**Fig. 2:**
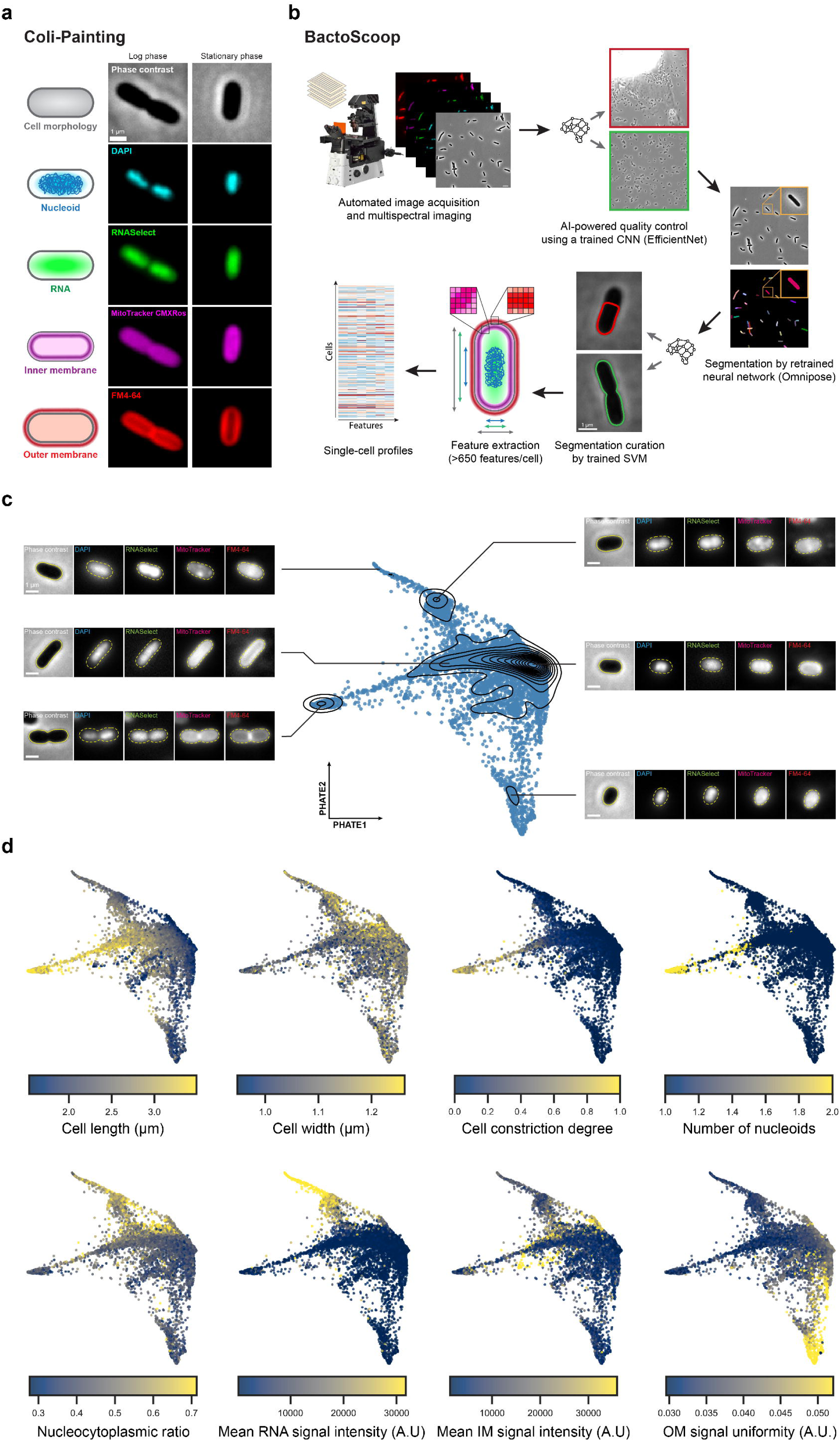
Coli-Painting and BactoScoop image-based profiling reveal population heterogeneity in stationary-phase *E. coli* cells. **a**, Overview of cellular structures targeted in Coli-Painting and the corresponding fluorescent dyes used for labeling. **b,** Schematic overview of the BactoScoop pipeline, illustrating the transformation of raw multispectral images into single-cell image-based profiles. **c,** Feature embedding of 652 extracted features across 12 WT replicates. Black contours indicate cell density, with representative cells shown from densely populated regions. **d,** The feature embedding shown in **c**, overlaid with features describing cell morphology, nucleoid, RNA, inner membrane, and outer membrane.

We first used this combined platform to further examine the single-cell level heterogeneity of *E. coli* in stationary phase. For this, wild-type *E. coli* cells were grown in M9glu, sampled in stationary phase (24 h after re-dilution), stained with the Coli-Painting dyes, and imaged on agarose pads. We chose this timepoint as it corresponds to early stationary phase, where cell biological feature behavior appeared to plateau upon completion of the transition from exponential growth (Fig. 1). We then applied BactoScoop to extract quantitative single-cell profiles. To obtain a systematic and reproducible analysis of phenotypic heterogeneity within stationary-phase populations, this approach was repeated across 12 independent replicates (Extended Data Fig. 2), yielding 39,576 single-cell level profiles in total (Dataset S2). The resulting embedding, based on a combination of 652 single-cell features, revealed the extensive cellular diversity that exists within these populations (Fig. 2c). Moreover, by projecting different individual features on the embedding, we could identify smaller subpopulations displaying specific properties related to different cell morphological aspects or other cellular components (Fig. 2c-d). This included previously noted subpopulations of longer cells displaying cell constriction and harboring two nucleoids, but also other groups of cells with specific RNA- or membrane-related properties (Fig. 2c-d).

### Genome-wide image-based profiling identifies deletion strains with deviating cell biological phenotypes in stationary phase

To interrogate the genetic basis of cell morphogenesis and subcellular organization in stationary-phase *E. coli* cells, we applied our image-based profiling approach to the *E. coli* Keio collection, a genome-wide library of single-gene deletion strains^47^. Mutants and wild-type strains were grown in M9glu, sampled during stationary phase (24 h after re-dilution), stained, imaged on large agarose pads, and subsequently analyzed using the BactoScoop pipeline. The final dataset comprises 15,860,005 single-cell profiles, each defined by 661 features, gathered from 3,703 deletion strains and 579 wild-type replicates, yielding an average of approx. 3,704 cells per strain or wild-type replicate. For all features, we identified strains displaying deviating behavior by calculating a population-level normalized score (*s*), which represents the extent to which a deletion strain deviates from the parent replicates (see Methods).

We first examined the cell morphological landscape of stationary-phase *E. coli* cells by plotting median cell length versus median cell width for each gene deletion strain and wild-type replicate (Fig. 3a). Wild-type replicates clustered tightly and most knockout mutants displayed cell morphologies similar to those of the wild type. However, a sizeable fraction of deletion strains (10.6%) displayed a deviating morphological phenotype (|*s*| ≥ 4 for median cell length, width, or both; see Methods), indicating a substantial genomic commitment to stationary-phase cell morphogenesis (Fig. 3a). This included several deletion strains with known cell division and/or cell separation defects during exponential growth, which carried over into stationary phase and gave rise to increased cell lengths in stationary phase as well (e.g., Δ*envC*^48,49^ and Δ*dedD^49,50^*). The same phenotypic persistence in stationary phase held true for several deletions of genes involved in envelope biogenesis or cell wall synthesis with reported increases in cell width during exponential growth (e.g., Δ*rodZ*, Δ*rffC,* Δ*rffH,* Δ*wecF*, and Δ*lpp^49,51–53^*^)^. At the same time, we also identified several morphological outliers containing deletions of genes that were not previously linked to cell morphogenesis, suggesting that these might serve as stationary-phase-specific determinants of cell morphology. Among these were genes of unknown function (so-called y-genes^54^; e.g., *yoaC*, *ydcS*, and *ysgA*), which thus likely exhibit stationary-phase-specific cellular roles, and genes that have previously been characterized, yet without being implicated in stationary-phase physiology. For example, RppH is an RNA pyrophosphohydrolase functioning as an mRNA-decapping enzyme that initiates RNA degradation by converting the 5’-terminal triphosphate or diphosphate of mRNAs to monophosphate, which triggers downstream processing by RNase E^55–57^. While deletion of *rppH* leads to an enhanced lifetime of hundreds of mRNAs^55,58^, our understanding of its physiological role remains rather limited. Although RppH activity has been implicated in stress resistance^59^ and envelope integrity^60^, the morphological deviations of stationary-phase Δ*rppH* cells (i.e., longer and wider cells) suggest that it might play a general role in mediating global mRNA turnover during transitions to non-growing cellular states, especially given that no morphological deviations were reported for a Δ*rppH* strain during exponential growth in our previous screens^49,51^. Across all strains, we did not identify a general relationship between median cell length and width (Spearman correlation (ρ) = 0.07), indicating that both are controlled independently in stationary phase, as we previously observed during exponential growth^49,51^.

**Fig. 3:**
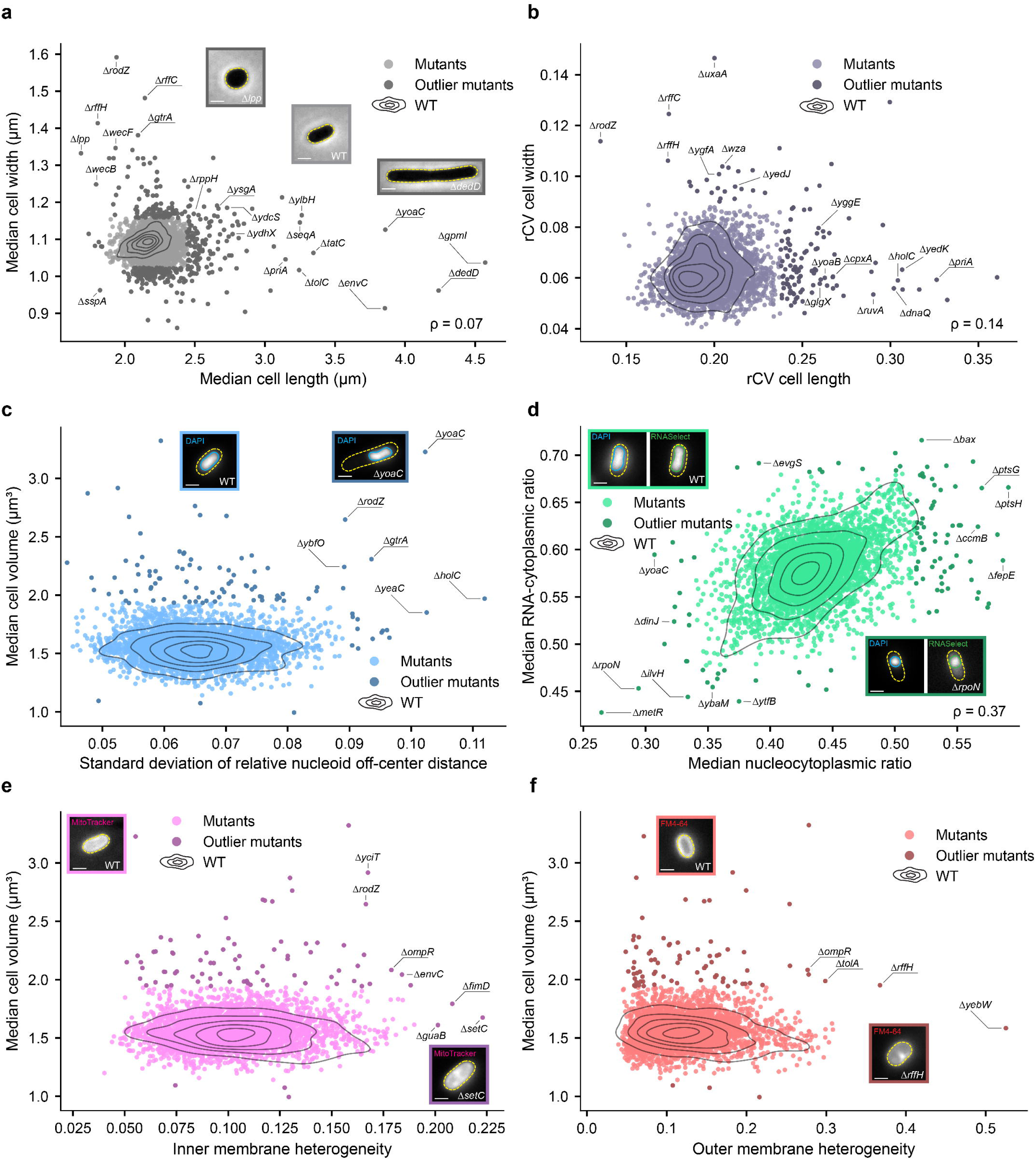
Examination of deviating gene deletion phenotypes in stationary phase. In all panels, wild-type (WT) populations are represented by black density contours corresponding to regions enclosing 10%, 30%, 50%, 70%, and 90% of the WT distribution. Colored dots represent deletion strains, with darker dots indicating strains deviating from the WT distribution (|*s*| ≥ 4). For all representative cells, the scale bar represents 1 µm. **a,** Scatter plot of median cell width versus median cell length. Spearman correlation coefficient (ρ) between both variables was calculated across all WT and deletion strains. Representative phase-contrast images of WT, Δ*lpp*, and Δ*dedD* cells are shown. **b,** Scatter plot of the robust coefficient of variation (rCV) in cell width versus rCV in cell length, including the corresponding ρ. **c,** Scatter plot of the median cell volume versus the standard deviation of relative nucleoid off-center distance. Representative nucleoid signal images of WT and Δ*yoaC* cells are shown. **d,** Scatter plot of the median RNA-cytoplasmic (RC) ratio versus the median nucleocytoplasmic (NC) ratio. Representative nucleoid and RNA signal images of WT and Δ*rpoN* cells are shown. **e,** Scatter plot of median cell volume versus inner membrane heterogeneity. Representative inner membrane signal images of WT and Δ*setC* cells are shown. **f,** Scatter plot of median cell volume versus outer membrane heterogeneity. Representative outer membrane signal images of WT and Δ*rffH* cells are shown.

We next assessed morphological heterogeneity within isogenic populations by examining the robust coefficient of variation (rCV; see Methods) of cell length and cell width across the deletion collection (Fig. 3b). Similar to the median cell length and width analysis, rCV cell length and rCV cell width were only weakly correlated (ρ = 0.14), indicating that strains typically exhibited increased heterogeneity in a single dimension. We observed elevated cell length variability (|*s*| ≥ 4) for several mutants of genes involved in DNA replication, repair, and genome maintenance (e.g., Δ*ruvA*^61–63^., Δ*holC^64^.*, Δ*dnaQ^65^.*, Δ*priA^63,66^.*, and Δ*yedK^67,68^.*). Likewise, we detected strains with increased cell width variability, including deletions of genes involved in cell envelope assembly and organization, such as *rffC^69^.*, *rffH^69^.*, *rodZ^70^.*, and *wza^71^.*. In addition, several outlier strains carried deletions of genes that had not previously been associated with morphological deviations nor increases in heterogeneity. This included both genes with characterized functions (e.g., Δ*uxaA^72^.* and Δ*cpxA^73^.*) and genes of unknown function (e.g., Δ*yggE* and Δy*edJ*). Moreover, several of the strains showing increased heterogeneity (e.g., Δ*glgX* and Δ*yoaB*) did not display a deviating median cell length or cell width, highlighting the strength of our single-cell level approach to identify variability within populations that would otherwise remain undetected.

Besides cell morphology, we identified outliers for features related to other major cellular components visualized by the dyes. This included deletion mutants displaying aberrant nucleoid positioning (Fig. 3c). In most strains, nucleoids were positioned centrally along a cell’s length axis, showing only small deviations from the cell center. However, several deletion strains exhibited a marked shift away from central nucleoid positioning, including genes of known function (e.g., Δ*rodZ^70^.* and Δ*holC^64^.*), but also genes of unknown function (e.g., Δ*yoaC*, Δ*ybfO*, and Δ*yeaC*). In contrast to exponential-phase cells, in which bulk RNA localization was largely homogeneously distributed, the RNASelect signal appeared more condensed and nucleoid-associated in stationary-phase cells (Fig. 2a, Fig. 3d). This was further underscored by the fact that the median RNA-cytoplasmic (RC) ratio (i.e., the area of the RNA signal divided by the cell area) was consistently and substantially smaller than 1 and that we observed a positive correlation (ρ = 0.37) between the RC ratio and the nucleocytoplasmic (NC) ratio (i.e., the nucleoid area divided by the cell area) (Fig. 3d). The latter indicates that both indeed tend to co-vary in terms of size. For both ratios, we identified deletion strains that displayed higher and lower values (e.g., Δ*metR*, Δ*rpoN*, and Δ*ilvH* with lower ratios and Δ*ptsG*, Δ*ptsH*, and Δ*bax* with higher ratios), but also deletion strains that no longer appeared to follow the overall positive trend (e.g., Δ*ybaM*, Δ*yoaC*, Δ*ytfB,* Δ*fepE,* Δ*ccmB*, Δ*evgS* and Δ*dinJ*). For the membrane stains, we identified several deletion strains that displayed different patterns than the uniform membrane staining observed in the wild type (Fig. 3e-f). For example, Δ*setC*, Δ*guaB*, and Δ*fimD* (for the inner membrane) and Δ*tolA*, Δ*rffH*, and Δ*yebW* (for the outer membrane) showed more heterogeneous patterns than the wild-type replicates (Fig. 3e-f).

To summarize, we obtained detailed phenotypic profiles for all sampled deletion strains (Dataset S3), including many strains containing deletions of genes previously implicated in stationary-phase physiology (Extended Data Fig. 3, Supplementary Text 1). Across all measured features, we identified outlier strains that broadly fall into two categories. First, deletions of genes of known function frequently exhibited additional and previously uncharacterized phenotypes in stationary phase, suggesting they may play a broader role in cellular physiology than currently recognized. Second, several deletions of genes of unknown function gave rise to distinct phenotypes in stationary phase, pointing towards potential cellular roles that are stationary-phase specific. As such, our dataset greatly expands the landscape of genes contributing to stationary-phase biology.

### Feature covariation reveals connections within stationary-phase cell architecture

To further probe potential relationships, or lack thereof, between population-level features that define the architecture of stationary-phase cells, we constructed a global feature correlation map for a selected subset of 13 directly interpretable features (Fig. 4a, Dataset S4). This analysis revealed expected relationships between several cell morphological features. For example, both median cell length and width correlated with median cell volume (ρ = 0.70 and ρ = 0.68, respectively; Fig. 4a). Consistent with the geometric coupling between average surface area and volume in rod-shaped bacteria^22,23,74^, median cell volume also correlated strongly with the median cell surface area (ρ = 0.95; Fig. 4a).

**Fig. 4:**
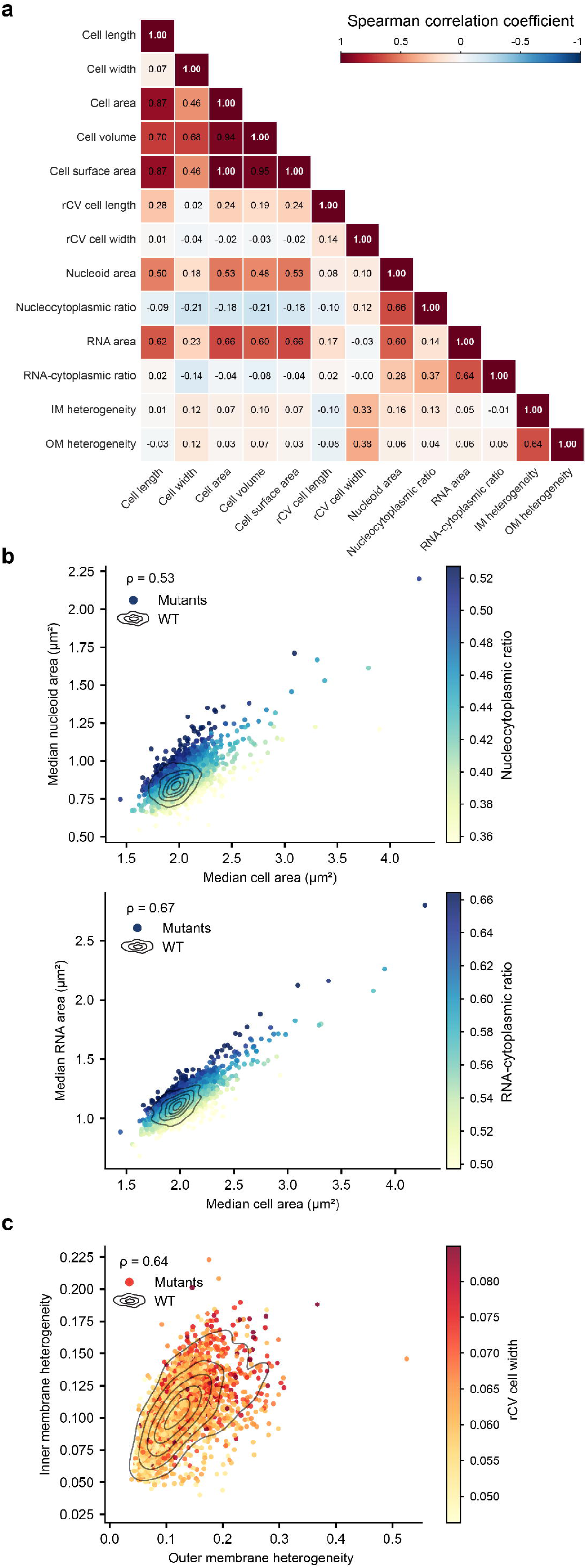
Genome-wide feature-feature correlations reveal cell-architectural connections in the stationary-phase cell. **a**, Heatmap showing pairwise correlations between 13 selected population-level features describing cell morphology and subcellular organization across the mutant and control populations. **b,** Scatter plots showing median nucleoid area versus median cell area colored by the NC ratio and median RNA area versus median cell area colored by RC ratio for wild-type (WT) and deletion strains. **c,** Scatter plots showing inner membrane heterogeneity versus outer membrane heterogeneity colored by the rCV cell width. In panels **b** and **c**, black density contours enclose 10%, 30%, 50%, 70%, and 90% of the WT distribution, with the Spearman correlation coefficient (ρ) indicating the association between both variables.

In line with the robustness of nucleoid size scaling in exponential phase^49,51,75^, the median nucleoid area of stationary-phase cells also scaled with their median cell area (Fig. 4a-b). Importantly, variation in the NC ratio was mostly driven by variation in nucleoid size, and thus chromosome compaction, instead of by variation in cell size (Extended Data Fig. 4a). We made similar observations for the RNA signal area (Fig. 4a-b), which scaled with cell and nucleoid area, and the RC ratio, where variation was mostly driven by variation in the RNA area (Extended Data Fig. 4b). In contrast, membrane-related features only showed a limited connection to other population-level features (Fig. 4a). Inner- and outer-membrane heterogeneity correlated positively (ρ = 0.64, Fig. 4c), suggesting that alterations in one of the membranes is often accompanied by alterations in the other. Moreover, increased heterogeneity in the membrane stainings was associated with more variable cell widths, as both inner and outer-membrane heterogeneity correlated positively with rCV cell width (ρ = 0.33 and 0.38, respectively). Together, these observations suggest a role for envelope integrity in maintaining a consistent cell width in stationary-phase cells.

### Profile correlations and global feature embeddings can provide functional context for gene deletions

To move from individual features to broader profile-level relationships, we next compared the phenotypic profiles of all strains across 412 selected features (see Methods). Pairwise Spearman correlations between strain profiles were used to identify mutants with similar stationary-phase morphologies and subcellular organization patterns (Fig. 5a, Dataset S5). In parallel, we constructed an embedding from the same feature space using Uniform Manifold Approximation and Projection (UMAP)^76^ (Fig. 5b), allowing the genome-wide collection to be visualized as a map in which neighboring strains share similar image-based profiles (Extended Data Fig. 5). To define relationships between strains more systematically, we performed a weighted morphological profiling network analysis (WMPNA; similar to a weighted gene co-expression network analysis for gene expression data^77,78^) on the image-based profiles (see Methods, Dataset S5). This analysis groups mutants with covarying profiles into modules and we projected the 20 largest modules onto the UMAP embedding (Fig. 5b). Most large modules formed localized neighborhoods on the map, indicating that the WMPNA module assignment captured coherent image-based profile similarities. In contrast, singletons, defined as genes belonging to modules with fewer than three strains, were mainly located in the wild-type-rich region of the embedding, suggesting that these strains did not display a strong shared deviation from the wild-type profile.

**Fig. 5:**
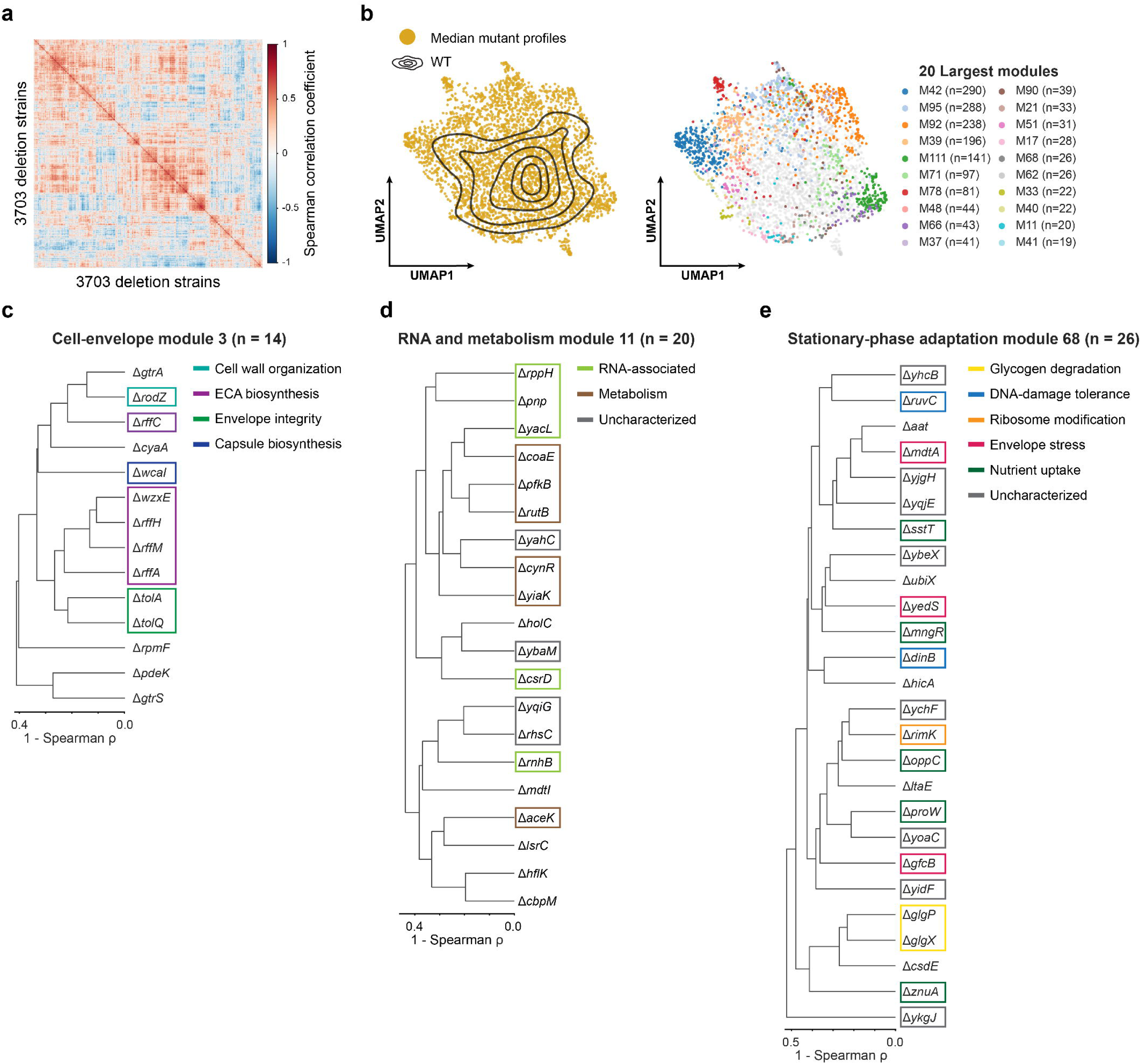
Profile correlations and feature embedding enable systematic analysis of functional neighborhoods. **a**, Heatmap showing the pairwise Spearman correlation coefficients between feature profiles of 3703 unique deletion strains. **b,** A visualization of the UMAP embedding of median mutant profiles described by 412 selected features, with black density contours corresponding to regions enclosing 10%, 25%, 50%, 75%, and 90% of the wild-type distribution and the UMAP embedding overlaid with the 20 largest WMPNA modules. **c,** Dendrogram showing the cell envelope module (WMPNA module 3; n = 14). **d**, Dendrogram showing the RNA and metabolism module (WMPNA module 11; n = 20). **e**, Dendrogram showing the stationary-phase adaptation module (WMPNA module 68; n = 26). In panels **c**, **d**, and **e**, the x-axis indicates 1 - Spearman correlation coefficient (ρ).

To illustrate the interpretive value of this map, we first focused on module 3, which grouped strains with a shared envelope-centered profile (Fig. 5c). The module included several genes involved in enterobacterial common antigen biosynthesis and transport (*rffH*, *rffM*, *rffA*, *rffC*, *wzxE*)^69,79,80^ and colanic acid biosynthesis (*wcaI*)^81^, linking the cluster to surface polysaccharide assembly. It also contained *tolA* and *tolQ*, which are part of the Tol-Pal system required for outer membrane integrity^82^, and *rodZ*, which connects the module to MreB-dependent cell-wall organization and synthesis^70^. This envelope-centered signature also extended to genes without canonical envelope annotations (e.g., *cyaA*, *rpmF*, *pdeK, gtrS,* and *gtrA*), suggesting they might play direct or indirect roles in envelope integrity. Second, we focused on module 11, containing the Δ*rppH* strain but also deletions of several other RNA-associated genes (Fig. 5d). Within this module, the profile of the Δ*rppH* strain correlated strongly with that of Δ*pnp* (Fig. 5d). PNP is another protein involved in mRNA turnover, functioning as a 3’ to 5’ exoribonuclease involved in general phosphorolytic mRNA degradation^83^. Further investigation of the phenotypic signatures of both deletion strains revealed deviating behavior that extends beyond cell morphology, with these cells not only being larger (both in terms of cell length and width) but also displaying smaller NC and RC ratios (Extended Data Fig. 6).

Beyond genes with previously characterized functions, this framework can also be used to explore the phenotypic context of genes of unknown function. We therefore examined module 68, which contained Δ*yoaC* (Fig. 5e), a strain characterized by an increased cell size, a low nucleocytoplasmic ratio, and asymmetrically positioned nucleoids (Fig. 3). Its profile neighborhood connected Δ*yoaC* to several stationary-phase-relevant functional groups, including glycogen degradation (Δ*glgP^84^.* and Δ*glgX^84,85^.*), DNA-damage tolerance and genome maintenance (Δ*dinB^86,87^.* and Δ*ruvC^88^.*), ribosome modification (Δ*rimK*)^89^, envelope stress and permeability-associated functions (Δ*mdtA^90,91^.*, Δ*yedS^92.^*, and Δ*gfcB^93,94^.*), and nutrient uptake (Δ*mngR^95^*, Δ*proW*^96–98^., Δ*sstT^99^*., Δ*oppC*^100,101^. and Δ*znuA^102^.*). The same module also contained several other uncharacterized or poorly characterized genes (Δ*yidF*, Δ*ybeX*, Δ*yqjE*, Δ*ykgJ*, Δ*yhcB,* Δ*ychF* and Δ*yjgH*), suggesting that their shared image-based profiles may reflect previously unrecognized links to stationary-phase cellular organization. More broadly, we also examined whether morphological diversity in stationary phase was associated with canonical stationary-phase regulatory programs. For this, we compiled gene-level annotations from the literature characterizing RpoS and ppGpp regulation in *E. coli* and assessed to what extent deletion strains of genes that are part of these regulons display a profile that deviates from the wild type (see Methods, Dataset S6). Across our collection, we found no clear association between deviating profiles and documented RpoS- and/or ppGpp-associations (Extended Data Fig. 7a). UMAP overlays also showed no clear segregation by regulatory program or direction of regulation (i.e., activated, repressed or mixed; Extended Data Fig. 7b).

### Deviating image-based profiles as gateways to uncover y-gene function

To further illustrate the potential of our approach to identify new genes that contribute to stationary-phase cell organization, we exploited the many deviating profiles of deletions of genes of unknown function. We selected a subset of 19 y-genes (*ybaB*, *ybaM*, *ychF*, *yeaC*, *yebV*, *yebW*, *yedJ*, *yedK*, *yggE*, *yggL*, *yhcB*, *yidF*, *yjgH*, *ynfA*, *yoaB*, *yoaC*, *yqjF*, *ysgA*, and *ytfB*) for further characterization, as we reasoned that the function of some of these genes has remained elusive because their roles emerge under conditions that have received limited attention, such as stationary phase. We first constructed C-terminal fluorescent protein fusions to each of them at their native locus, which enabled us to determine growth-phase-dependent changes in protein levels and intracellular localization patterns (Extended Data Fig. 8). Quantification of average single-cell fluorescence revealed a stationary-phase increase for 8 fusions (Fig. 6a, Dataset S7), and even a decrease for a single fusion (i.e., YeaC-msfGFP), underscoring the strength of our approach in identifying relevant gene candidates. Comparison of our measurements to published transcriptomic^6,7,103–105^, proteomic^8,15^, and mRNA half-life^104^ datasets revealed a mixed picture (Fig. 6b, Dataset S8). While increases in fluorescence were consistent with increases in transcript abundance and/or protein levels for some genes (e.g., *yebV*, *yjgH*, and *yggE*), other genes displayed discrepancies between measurements (often also across published datasets). For example, YbaB-msfGFP levels were increased in our setup despite displaying decreased RNA and protein abundance in all available datasets (Fig. 6b). Also, all nine candidates with mRNA half-life measurements exhibited increased transcript stability in stationary phase, including genes of which the transcript abundance decreased (e.g., *yoaB*) and genes of which the fluorescent protein fusion displayed lower intensities (e.g., *yeaC*). Together, these findings revealed both convergent and divergent patterns of growth-phase-dependent regulation across different setups and measurement approaches.

**Fig. 6:**
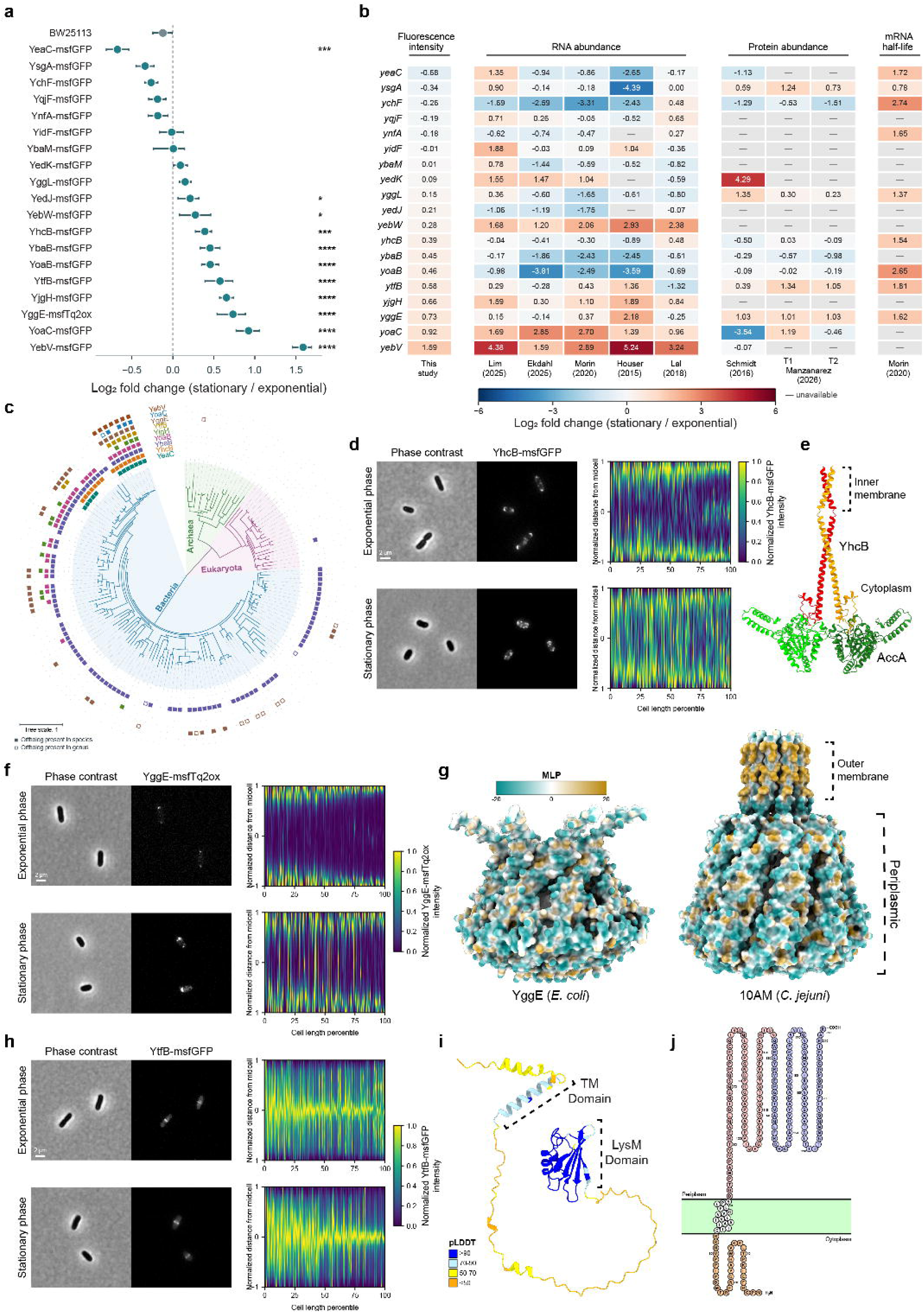
Characterization of genes of unknown function. **a**, Forest plot showing changes in fluorescence intensity from exponential to stationary phase across the indicated fusions. Points show log₂ fold changes calculated as log₂(stationary/exponential). Three independent replicates were analyzed per strain and phase, except for YjgH-msfGFP (n = 2). Error bars show 95% bootstrap confidence intervals obtained by resampling replicate means. Asterisks indicate differences in the phase-dependent change relative to the control, tested using two-sided contrasts from a strain × phase linear model with Holm correction. Adjusted *p*: * < 0.05, ** < 0.01, *** < 0.001, **** < 0.0001. **b**, Heatmap comparing fluorescence changes with published RNA abundance, protein abundance and mRNA half-life log₂ fold changes. Manzanarez T1 and T2 denote 6 and 24 h after glucose exhaustion, respectively. **c**, Orthologs of the indicated proteins, identified using eggNOG 5.0, were mapped onto the tree of life. Filled squares indicate supported presence in the displayed species, open squares indicate presence only in another sampled species of the same genus. No square indicates not detected or unresolved. Species names are provided in Extended Data Fig. 9b. **d**, Left, representative phase-contrast and epifluorescence images of the BW25113 *yhcB::yhcB-msfgfp* strain in exponential phase (top row) and stationary phase (bottom row). Right, demographs of the YhcB-msfGFP signal in exponential phase (top) and stationary phase (bottom). **e**, AlphaFold 3-predicted structure of a YhcB homodimer in complex with an AccA homodimer, the α-subunit of acetyl-CoA carboxylase, illustrating the predicted interaction between the two proteins. **f**, Left, representative phase-contrast and epifluorescence images of the BW25113 *yggE::yggE-msftq2ox* strain in exponential phase (top row) and stationary phase (bottom row). Right, demographs of the YggE-msfTq2ox signal in exponential phase (top) and stationary phase (bottom). **g**, AlphaFold 3-predicted structure of *E. coli* YggE, surface colored according to Molecular Lipophilicity Potential (MLP). In addition, the experimentally determined structure of the SIMPL-like protein, which forms a nonameric complex, from *Campylobacter jejuni* (PDB: 10AM), colored by MLP. The transmembrane and periplasmic domains are highlighted accordingly. **h**, Left, representative phase-contrast and epifluorescence images of the BW25113 *ytfB::ytfB-msfgfp* strain in exponential phase (top row) and stationary phase (bottom row). Right, demographs of the YtfB-msfGFP signal in exponential phase (top) and stationary phase (bottom). **i**, AlphaFold 3-predicted structure of YtfB, colored by pLDDT. The predicted transmembrane (TM) and the OapA (LysM-like) domain are highlighted and linked by a 79-residue region with low prediction confidence. **j**, Topology diagram of YtfB, illustrating cytoplasmic, inner membrane, and periplasmic domains.

To further assess the biological significance of the nine gene fusions displaying significant changes in fluorescence levels in our setup (Fig. 6a), we also investigated their evolutionary conservation. Within *E. coli*, across 136,051 genomes, the conservation of these genes exceeded 99% except for *yjgH* (87.439%) and *ytfB* (91.229%) (Extended Data Fig. 9a). More broadly, across the tree of life, only *yggE* and *ybaB* displayed a somewhat extended distribution (Fig. 6c; Dataset S9), with others being more constrained to specific bacterial classes (Extended Data Fig. 9b). AlphaFold 3 prediction also revealed a broad diversity of protein structures encoded by these y-genes (Extended Data Fig. 10a).

Subsequently, we examined three candidates exhibiting a distinct subcellular localization pattern in more detail. YhcB-msfGFP formed discrete fluorescent foci that preferentially concentrated at the poles and midcell during exponential growth, but became more widely distributed throughout the cell during stationary phase (Fig. 6d). Previous studies have implicated YhcB in cell division, peptidoglycan synthesis, and lipopolysaccharide biosynthesis^106–109^, with the most recent ones forwarding a role for YhcB in inhibiting fatty acid biosynthesis in a growth-phase-specific way by sequestering AccA and inhibiting acetyl-coenzyme A (CoA) carboxylase (ACC) complex formation in early stationary phase^110,111^. Our findings support the stationary-phase specific role of YhcB, with the redistribution in protein localization (from polar and midcell to distributed across the entire cell; Fig. 6d) likely reflecting an additional layer of regulation. Moreover, AlphaFold 3 modelling supported the formation of a YhcB_2_–AccA_2_ complex (Fig. 6e), whereas modelling YhcB in association with the assembled acetyl-CoA carboxylase (ACC) complex suggested that this arrangement is structurally unfavorable and not compatible with YhcB inserting into the membrane (Extended Data Fig. 10b). As such, these predictions are consistent with a mechanism in which YhcB sequesters AccA and prevents its incorporation into the active ACC complex (Fig. 6e; Extended Data Fig. 10b).

YggE, a conserved but poorly characterized protein that has been implicated in protection against oxidative stress^112,113^, exhibited polar localization during both exponential and stationary phase, with increased fluorescence intensity and a broader polar signal during stationary phase (Fig. 6a, f). YggE displays structural homology to the BP26 protein of unknown function from *Brucella abortus*^114^ (PDB: 4HVZ; Extended Data Fig. 10c) and the SIMPL-like protein Cj0034c from *Campylobacter jejuni*, which forms a nonameric channel-like structure (PDB: 10AM)^115^. Unlike the *C. jejuni* protein, a YggE complex lacks the hydrophobic segment likely required for membrane insertion (Fig. 6g). The cage-like complex creates a continuous internal lumen, 73.82 Å long and up to 78.75 Å wide (Extended Data Fig. 10d). Similar to BP26 and Cj0034c, the exact function of YggE remains to be resolved, but our data suggest a spatially organized role for this complex that may become particularly relevant in stationary phase.

Another protein displaying a distinct localization pattern and an increased concentration in stationary phase was YtfB (Fig. 6a, h). Previous studies have identified YtfB as an inner membrane cell division protein with an N-terminal transmembrane helix and a C-terminal OapA domain that binds peptidoglycan^116^ (Fig. 6i). YtfB is recruited early to the divisome at the future division site in an FtsZ-dependent manner in exponentially growing cells^116^. Our data, using production of a YtfB-msfGFP fusion from its native locus instead of relying on plasmid-based production, show that this midcell divisome-association persists in stationary phase, with increased levels of YtfB (Fig. 6h). Through its domains, YtfB connects the divisome in the inner membrane with the peptidoglycan layer (Fig. 6i-j). While this bridge function has been proposed to help coordinate early divisome assembly with later stages of septal peptidoglycan remodeling, its relevance during stationary phase remains to be established.

## DISCUSSION

Stationary phase is commonly defined by the absence of net population growth, but this captures only the population-level outcome of a broader cellular transition. As *E. coli* approaches growth arrest, morphology, chromosome organization and replication, and cell division change progressively, with individual cells occupying overlapping states along the transition (Fig. 1). This diversity persists even after net population growth is halted, with populations that span a broad phenotypic space encompassing differences in morphology, nucleoid organization, RNA and membrane properties (Fig. 2). Genome-wide perturbation further demonstrated the large genomic contribution to different aspects of this phenotypic space (Fig. 3), which we exploited to identify dependencies between cellular features (Fig. 4) and functional connections between genes (Fig. 5). We also leveraged such connections to provide insight into the contribution and potential role of several genes of unknown function to stationary-phase cell morphogenesis and subcellular organization (Fig. 6). Together, our analysis highlights stationary phase as a heterogeneous and time-variant phase of which the properties behave as complex phenotypic traits shaped by the combined effects of many different genes.

Our study identified a wealth of new genes associated with stationary-phase physiology, including both genes of known function with previously underappreciated contributions to stationary phase and genes of unknown function with potential stationary-phase-specific roles. Examples of the former include *rppH* and *pnp*, of which the stationary-phase-specific deviating behavior of their corresponding deletion strains suggest an important role for mRNA degradation and turnover upon transitioning to stationary phase, while examples of the latter include all the y-genes we further characterized. All of this was made possible via the Coli-Painting assay and BactoScoop image-based profiling pipeline we developed, which substantially expands the phenotypic space in which the effect of a gene deletion can be detected. This contrasts the approach of many other systematic screens, which often reduce the consequence of a perturbation to a limited set of phenotypes, most commonly growth or fitness. Instead, our pipeline simultaneously reports on many aspects of cellular physiology, which proved particularly informative in the growth-limiting conditions of our screen. Moreover, our pipeline generates higher-dimensional profiles in higher throughput than previous image-based approaches used to interrogate the genetic basis of cell morphogenesis and cell cycle progression during exponential growth in *E. coli^49,51,117,118^*. Furthermore, we expect our approach to be readily transferable to other environments and growth conditions, opening up new avenues for quantitative cell biological studies of *E. coli* across conditions and in more realistic suboptimal environments (e.g., in the presence of different stressors).

The power of our approach also lies in the combination of features rather than in their number alone. While measurements of individual features can identify conspicuous phenotypes, many genetic perturbations produce coordinated changes across cell morphology and subcellular organization that are difficult to interpret in isolation. Integrating these measurements into profiles revealed similarities between mutants and recovered functionally coherent neighborhoods (modules) in phenotype space (Fig. 5). For example, perturbations affecting surface-polysaccharide synthesis, Tol-Pal function and cell-wall organization converged within an envelope-centered module despite acting at distinct steps and locations of envelope biogenesis. Such convergence suggests that high-dimensional profiles capture shared cellular consequences of genetic perturbations rather than simply extreme values of specific individual features. On the other hand, it is important to note that phenotypic similarity of genetic perturbations does not imply members of a module to be part of a common molecular pathway. Instead, it provides an indication that these perturbations lead to similar phenotypic outcomes, which can guide future functional characterization experiments.

Despite extensive phenotypic variability across thousands of genetic perturbations, several aspects of stationary-phase cell architecture remained closely coupled. Inner- and outer-membrane heterogeneity correlated with each other and with cell-width variability, suggesting a connection between envelope organization and morphological stability. Nucleoid size also correlated with cell size. This nucleoid size scaling was somewhat less strong than in exponentially growing cells (ρ = 0.53 in stationary phase versus ρ ≥ 0.71 during exponential growth^49,51,75^), likely due to the diminished presence of DNA replication and the fact that the majority of cells carry just a single copy of their chromosome at this stage. However, the observed scaling does imply that smaller stationary-phase cells package their single chromosome into a more compact and dense nucleoid than larger cells. Similarly, the RNA-stained area scaled with cell and nucleoid size. This relationship would have remained undetected without the multiplexed staining approach, highlighting the value of simultaneously resolving multiple subcellular compartments.

Although *E. coli* is among the most extensively studied organisms, a substantial fraction of its genes remains poorly characterized. These unannotated genes are typically referred to as y-genes, stemming from their primary names starting with the letter y^54,119^. Part of the explanation for this sequence-to-function gap lies in the fact that the function of these y-genes may only become apparent under specific physiological conditions and that the proper condition has simply not been probed yet. This highlights the importance of investigating gene deletion phenotypes across different environments, beyond exponential growth. Our genome-wide microscopy screen further underscores this by identifying many deletions of poorly characterized genes that are associated with stationary-phase-specific alterations in cell morphology and subcellular organization. Combining these deviating profiles with subsequent examination of protein concentration, protein localization, structural properties, and phylogenetic conservation can then serve as gateways for uncovering protein function.

In conclusion, our study shows that stationary phase is not simply the stage at which population growth stops, but a distinct and heterogeneous state of cellular (re)organization. This restructuring develops progressively, varies substantially between individual cells, and is dependent on the function of a large fraction of the non-essential genome. Coli-Painting and BactoScoop made this restructuring measurable in the form of quantitative phenotypic features, providing a detailed phenotypic reference for all included deletion strains. Together our approach highlights the power of bacterial image-based profiling to uncover gene function and distinguish physiological states that would otherwise remain unresolved.

## MATERIALS & METHODS

### Experimental procedures

#### Strains and growth conditions

Bacterial strains and primers used in this study are listed in Tables S1 and S2, respectively. Strains were cultured in M9 medium (6 g/l Na_2_HPO_4_, 3 g/l KH_2_PO_4_, 0.5 g/l NaCl, 1 g/l NH_4_Cl, 2 mM MgSO_4_, 0.1 mM CaCl_2_), supplemented with 0.2% (w/v) glucose (M9glu), or in LB (5 g/l yeast extract, 10 g/l tryptone, 5 g/l NaCl). For all experiments, pre-cultures of each strain were grown to stationary phase in the appropriate growth medium (200 µl per well) in 96-well F-bottom plates (Greiner BIO-ONE) covered with sterile lids (Greiner BIO-ONE) at 37°C under well-aerated conditions (200 rpm orbital shaking). Pre-cultures were then diluted 1:10,000 into fresh medium (200 µl per well) and allowed to grow under the same conditions. Cultures were subsequently sampled for microscopy at indicated times.

#### Strain construction

Mutant strains producing C-terminal translational fluorescent fusions to y-genes of interest were constructed in *E. coli* BW25113, the parent strain of the Keio collection, equipped with the pKD46 plasmid^120^. Detailed construction procedures for each mutant strain are given in Table S1. To construct C-terminal translational fusions of the gene of interest with *msfgfp*, the *msfgfp-frt-nptI-frt* cassette was amplified by PCR from plasmid pDHL1029*^121^*, using primers with ±50 bp tails complementary to the target chromosomal region, replacing the stop codon of the gene of interest with the *msfgfp-frt-nptI-frt* cassette. For most strains, the kanamycin resistance gene (*nptI*) was excised by the Flp site-specific recombinase via transient equipment of the strains with plasmid pCP20^122^. All constructed mutants were initially confirmed by PCR with primer pairs binding outside of the region where homologous recombination occurred, and further verified by Sanger sequencing (Eurofins). Sequencing results were analyzed in Benchling by aligning the sequencing reads to the *in-silico*-designed chromosomal sequences using default parameters (Multisequence alignments, MAFFT v7^123^).

#### Mapping single-cell behavior of *E. coli* populations along their growth curve

Precultures of *E. coli* OZA001^43^ cells, producing GFP-DnaN, were grown in 96 well plates for 24 h in 200 µl M9glu (37°C, 200 rpm), diluted 1:10,000 into fresh medium (200 µl), and sampled at 13 timepoints (9, 10, 12, 14, 16, 20, 22, 23, 24, 26, 28, 46, and 50 h after re-dilution) spanning most of the growth curve (Fig. 1a). To visualize the nucleoid, which is the intracellular region that contains the chromosomal DNA, cells were incubated in their growth medium with DAPI (Merck Life Science, D9542) at a final concentration of 1 µg/ml for 15 min prior to sampling for microscopy. For each sample, 0.5 µl of culture was spotted on an agarose pad and covered with a glass coverslip. This time-course experiment was also performed in LB medium using a similar methodology. LB cultures were sampled 3, 4, 6, 8, 12, 16, 18, 20, and 30 h after re-dilution.

#### Coli-Painting

To perform Coli-Painting, cultures of cells were first stained with SYTO™ RNASelect™ Green Fluorescent Cell Stain (final concentration of 500 nM; Thermo Fisher Scientific, S32703) and MitoTracker™ Red CMXRos (final concentration of 1 µg/ml; Thermo Fisher Scientific, M7512), and incubated for 30 min at 37 °C with shaking (200 rpm) in the dark. Cells were subsequently fixed by the addition of methanol-free formaldehyde (16% stock; Thermo Fisher Scientific, 28906) to a final concentration of 4% (v/v) and incubated for 30 min at 4 °C in the dark. Fixed cells were washed twice in M9 buffer by centrifugation at 2000 rpm for 10 min at room temperature. Cells were then stained with DAPI (final concentration of 1 µg/ml; Merck Life Science, D9542) and FM™ 4-64FX membrane dye (final concentration of 1 µg/ml; Thermo Fisher Scientific, F34653). Samples were incubated for 30 min at room temperature in the dark without shaking, after which the samples were imaged.

#### Genome-wide microscopy screen

The Keio collection mutants were transferred from frozen stocks stored at −80°C in 25% glycerol (96-well format) onto LB agar supplemented with kanamycin (50 μg/ml) using a 96-pin replica plater. Plates were incubated overnight at 37°C. Pre-cultures were generated by transferring colonies to 96-well F-bottom plates with 200 µl M9glu and grown for 24 h at 37°C with shaking at 200 rpm. These precultures were then diluted 1:10,000 in 200 µl M9glu and grown for 24 h. From each stationary-phase culture, 60 µl was transferred to a new 96-well plate and subsequently subjected to the Coli-Painting protocol before imaging.

#### Fluorescent y-gene fusions

To investigate the up- or down-regulation and subcellular localization of selected genes of unknown function during the transition from exponential to stationary phase, fluorescent protein fusions were generated in *E. coli* BW25113. The selected strains produced YeaC-msfGFP, YsgA-msfGFP, YchF-msfGFP, YqjF-msfGFP, YnfA-msfGFP, YidF-msfGFP, YbaM-msfGFP, YedK-msfGFP, YggL-msfGFP, YedJ-msfGFP, YebW-msfGFP, YhcB-msfGFP, YbaB-msfGFP, YoaB-msfGFP, YtfB-msfGFP, YjgH-msfGFP, YggE-msfTq2ox, YoaC-msfGFP, and YebV-msfGFP (Table S1). Colonies were inoculated into 96-well F-bottom plates containing 200 µl M9 glucose medium and incubated for 24 h at 37 °C with shaking at 200 rpm. Following a 1:10,000 dilution into fresh medium, cultures were sampled at 12 h and 24 h to capture exponential and stationary growth phases, respectively. For microscopy, 0.5 µl of each sample was spotted on an agarose pad, covered with a glass coverslip, and imaged within 5-10 min of preparation.

#### Microscopy

For all experiments excluding the genome-wide screen, cells were imaged on small (1.5%) agarose pads (∼25mm × 25mm) made by dissolving UltraPure Agarose (Invitrogen, Thermo Fisher Scientific, USA) in M9 buffer. For each sample, 0.5 µl of cell culture was spotted on the pad, which was then covered with a glass coverslip (Fisherbrand Glass Square Coverslips, Fisher Scientific, Belgium). Per sample, ∼20 positions were imaged to ensure capture of a sufficient number of cells. For the genome-wide screen, a high-throughput automated imaging workflow was implemented. Cells were imaged on large (3%) agarose pads prepared by dissolving UltraPure Agarose (Invitrogen, Thermo Fisher Scientific, USA) in M9 buffer (25 ml). Following heating, the dissolved agarose was poured into the lid of a 96-well plate with individual condensation rings (9 mm depth) and allowed to solidify. The resulting agarose slab was excised using a scalpel to generate a pad corresponding to half of a 96-well plate (48 wells). For each sample, 4 µl of cell culture was spotted onto the agarose pad in an 8 × 6 grid layout, after which the pad was transferred to a single-well glass-bottom imaging plate (Cellvis, USA) for microscopy. To enable fully automated imaging of the genome-wide screen, NIS-Elements JOBS (Nikon) was used to implement a scripted acquisition workflow that systematically imaged 20 positions for all 48 samples across the agarose pad.

Phase-contrast and epifluorescence imaging was performed on a Nikon Eclipse Ti2-E inverted epifluorescence microscope controlled by NIS-Elements AR software. The microscope was equipped with a Plan Apo λ 100× DM oil immersion objective (NA = 1.45), a Ph3 condenser annulus, and a Kinetix A22J723021 sCMOS camera (Teledyne Photometrics, USA). Fluorescence excitation was provided by a pE-800 LED light source (CoolLED, UK) with eight independently controlled LED channels. Narrowband excitation and emission were achieved using BrightLine Pinkel filter sets (penta-band set: LED-DA/FI/TR/Cy5/Cy7-5X-A; Semrock, IDEX Health and Science, USA) in combination with OptoSpin filter wheels (Cairn GmbH, Germany). The following optical configuration was used to acquire fluorescent images: GFP (excitation [CoolLED]: 470 nm, excitation filter: FF01-474/27, dichroic FF409/493/573/652/759-Di01, emitter filter cube: FF01-432/515/595/681/809, emitter filter wheel: FF01-515/30), DAPI (excitation [CoolLED]: 400 nm, excitation filter: FF01-378/52, dichroic FF409/493/573/652/759-Di01, emitter filter cube: FF01-432/515/595/681/809, emitter filter wheel: FF01-432/36), mCherry (excitation [CoolLED]: 580 nm, excitation filter: FF01-578/21, dichroic FF459/526/596-Di01, emitter filter cube: FF01-475/543/600, emitter filter wheel: FF01-641/75), FM4-64 (excitation [CoolLED]: 500 nm, excitation filter: FF01-509/22, dichroic FF459/526/596-Di01, emitter filter cube: FF01-475/543/600, emitter filter wheel: FF01-809/81).

### Image processing and statistical analysis

#### BactoScoop image-based profiling

##### Image-level quality control

Low-quality images were removed prior to downstream analysis using a convolutional neural network classifier trained to distinguish usable from non-informative fields of view. The classifier was based on the EfficientNet-B0^124^ architecture and was trained on phase-contrast images manually annotated as either acceptable or poor quality. The latter included images affected by blurring, overcrowding, saturation artifacts, or absence of cells. The training dataset consisted of 327 high-quality and 210 low-quality images. Images were resized to 224 × 224 pixels, and the model was trained using a batch size of 8 with an 80:20 training-validation split and a learning rate of 1 × 10^-4^. Early stopping was applied after 5 epochs to avoid overfitting. Model performance was evaluated on an independent test set containing 34 high-quality and 25 low-quality images, achieving an accuracy of 0.96. After classification of the full dataset originating from our genome-wide screen, images predicted as poor quality were visually inspected to identify potential false negatives and to ensure that biologically relevant cells were not excluded from downstream analysis.

##### Image analysis and feature extraction

For image analysis, we developed a custom Python-based software package that enables high-throughput bacterial single-cell analysis. This package integrates segmentation, control over whether segmented cells are dividing or already divided, internal coordinate reconstruction, and feature extraction into a unified workflow optimized for multichannel microscopy datasets. Cells were segmented from phase-contrast images using Omnipose^125^ (v1.0.6), a deep learning method designed to accurately delineate bacterial cell boundaries across diverse morphologies. Throughout this study, the pretrained bacterial model *bact_phase_omni* was used for segmentation. As the presence of adjacent cells can affect the extraction of cellular features, highly connected segmentation masks were removed using neighborhood-based filtering to reduce clumped detections (e.g., 4 or more neighbors) in densely populated fields of view. To correct instances where Omnipose oversegmented individual cells, segmented objects were merged when their pole-to-pole spacing fell below a defined distance threshold (≤ 4 pixels). Each obtained cell mask was transformed into an internal coordinate representation (“mesh”) derived from the cell contour and a midline. The midline was obtained by skeletonizing the mask, smoothing the resulting skeleton, and extending it to the cell poles. Lines perpendicular to the midline were intersected with the smoothed cell contour, yielding paired boundary points that defined successive cross-sections of the mesh. The cell mesh serves as the basis for the quantification of all morphological and fluorescence-derived features. Meshes were subsequently evaluated for constriction along their longitudinal axis to identify potential division events or remaining segmentation artifacts. Cells exceeding a configurable constriction degree threshold (cell_constriction_degree_width_axial > 0.5) were split at the constriction minimum. Cell constriction degree was calculated in similar fashion as in Oufti^126^, from the combined contour geometry and an intensity-derived signal obtained from inverted phase-contrast images. Together, these complementary joining and splitting operations provided consistent control over whether segmented objects are interpreted as single cells or as dividing cells, minimizing propagation of segmentation artifacts into downstream quantitative measurements.

Mesh quality control was then performed using a supervised support vector machine (SVM) classifier to distinguish high-quality from poor-quality cell detections, as before^127^. The classifier for this study was trained on a curated dataset comprising 74,194 manually labeled cells, with a positive-to-negative label ratio of 0.63:0.37. Training was performed using an 80:20 train-test split and feature dimensionality reduction via a Nyström approximation (100 components) combined with a radial basis function (RBF) kernel (Nyström γ = 0.01). The SVM was trained with regularization parameter C = 1000 and kernel coefficient γ = 1. Model performance was evaluated using 5-fold cross-validation, yielding a cross-validation accuracy of 0.951. Evaluation on the independent test set resulted in an accuracy of 0.950 and area under the receiver operating characteristic curve (AUC) of 0.990, with precision of 0.917, recall of 0.951, specificity of 0.949, and F1-score of 0.934. Cells classified as unfit for single-cell analyses were excluded from downstream analyses to improve robustness of feature extraction.

Subcellular fluorescent objects were detected using Laplacian-of-Gaussian-based blob detection (default σ = 3), followed by morphological refinement and assignment of detected objects to individual cell interiors based on spatial overlap criteria. Using the curated meshes, single-cell features were subsequently extracted from both cell shape and associated fluorescence signals, capturing >650 descriptors related to cellular morphology and subcellular organization. These include geometric, spatial, intensity, relationships between fluorescence channels, and object-based measurements that quantify cellular structure and intracellular signal distributions. Definitions, naming conventions, and measurement units for all extracted features are provided in the corresponding datasets (Dataset S2-3).

The entire BactoScoop pipeline is implemented in Python (v3.10) and supports parallel processing of microscopy datasets. Detailed installation instructions, documentation, and step-by-step workflow examples enabling reproducible execution of the pipeline are available in the project GitHub repository: https://github.com/Govers-Lab/bactoscoop.

#### Single-cell image-based profiles from the *E. coli* GFP-DnaN time-course experiments

##### Dataset generation, quality control, and population-level analysis

*E. coli* cells producing GFP-DnaN were grown in M9 glucose medium (M9glu) and sampled at 13 timepoints: 9, 10, 12, 14, 16, 20, 22, 23, 24, 26, 28, 46, and 50 h. Following feature extraction with BactoScoop, the combined dataset comprised 66,052 cells from three biological replicates. Extreme size outliers were identified separately within each replicate using absolute z-scores for cell_volume, cell_length, and cell_area, with a threshold of 8 standard deviations. Cells were subsequently retained only when the NC ratio (C2_NC_ratio) was within a physiologically possible range (0.05-1.1). The final M9glu dataset contained 64,621 single-cell profiles.

Morphological features were summarized over time by first calculating the mean value for each replicate at each timepoint and then averaging across replicates. Standard deviations across biological replicates were used as error estimates. The analyzed features included cell length, volume, aspect ratio, surface-area-to-volume ratio, width, area, and surface area. Fraction-based features were calculated independently for each replicate and timepoint by dividing the number of cells satisfying a specified condition by the total number of cells in that replicate-time group. These included the fractions of cells with no more than one nucleoid object (C2_object_number = 1), a low DnaN-to-cytoplasmic ratio (C3_NC_ratio ≤ 0.4), and detectable constriction (cell_constriction_degree_width_axial ≥ 0.1). Mean fractions and standard deviations were then calculated across replicates at each timepoint. Sigmoid curves were fitted to visualize temporal trends.

An analogous GFP-DnaN time-course experiment was performed in LB medium and is presented in Extended Data Fig. 1d-e. The LB dataset was subjected to the same feature extraction, quality-control, and population-level aggregation procedures. No PHATE analysis was performed for the LB experiment.

##### Time-course single-cell PHATE embedding

To visualize changes in single-cell state across the M9glu growth curve, PHATE was applied to cells collected at 12, 14, 20, 24, and 46 h. The input contained 13 cell-morphology and nucleoid-associated features (Dataset S1). Each timepoint was randomly downsampled to 2,922 cells to ensure balanced representation, yielding 14,610 cells in total. Features were standardized before dimensionality reduction. PHATE was run with knn = 10 and random_state = 42, producing a two-dimensional embedding used to visualize temporal changes in single-cell morphology.

#### Single-cell image-based profiles of wild-type *E. coli*

##### Single-cell profile quality control and balanced sampling

Single-cell image-based profiles of wild-type *E. coli* were analyzed to visualize population heterogeneity across 12 selected control wells. Cells were retained only when the outer membrane-to-cytoplasmic ratio ranged from 0.8 to 1.3, thereby excluding cells affected by detection errors. After filtering, 55,984 cells remained. To prevent wells containing larger numbers of cells from disproportionately influencing the embedding, each well was randomly downsampled without replacement to match the smallest retained well. This yielded 3,298 cells per well and 39,576 cells in total. Sampling was performed with a fixed random seed to ensure reproducibility.

##### PHATE embedding construction and feature visualization

All non-metadata columns (661) were initially considered as candidate morphological features. Features with more than 50% missing values in the downsampled dataset were excluded, and remaining missing values were imputed using the corresponding feature median. In total, 198,280 values were imputed, representing 0.77% of the PHATE input matrix. The final input matrix comprised 39,576 cells and 652 morphological features, which were standardized to zero mean and unit variance before dimensionality reduction (Dataset S2).

Single-cell heterogeneity was visualized in two dimensions using PHATE with n_components = 2, knn = 20, decay = 50, and automatic diffusion-time selection (t = “auto”). PHATE selected a diffusion time (t = 22), and the resulting coordinates were used for all downstream single-cell visualizations. Features were overlaid onto the fixed PHATE embedding using color limits defined by the 1^st^ and 99^th^ percentiles.

#### Genome-wide image-based profiles

##### Profile aggregation

Single-cell morphological profiles were processed in Python using pandas, NumPy, and pycytominer^128^. The dataset comprised 15,860,005 single-cell profiles described by 661 features. Feature-name prefixes indicated the imaging channel from which each measurement was derived: C1 corresponds to the phase-contrast images, C2 the outer-membrane channel, C3 the inner-membrane channel, C4 the RNA channel, and C5 the nucleoid channel.

Single-cell profiles were aggregated to well-level profiles by taking the median of each feature within each plate-well group, yielding 4,331 well-level profiles. For feature *f* in well *w* and plate *p*, the aggregated value was calculated as:

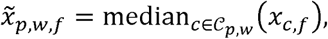

where *c* denotes an individual cell and *C_p,w_* is the set of cells assigned to plate *p* and well *w*. Visually confirmed low-quality and low cell-count (n < 50) samples were excluded before downstream processing, leaving 4,282 well-level profiles.

##### Plate-median rescaling

To reduce plate-specific shifts, each feature was rescaled so that its plate-wise median matched the corresponding median across the full dataset. Non-negative features were adjusted multiplicatively in log space using ε = 10^-6^:

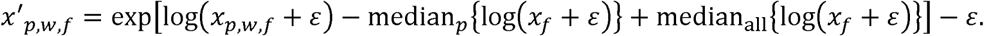

Signed features were adjusted additively:

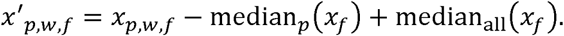

##### Calculation of population heterogeneity features

Additional descriptors of within-well single-cell heterogeneity were calculated from the original single-cell distributions and merged with the well-level table. Coefficients of variation were calculated for 20 fluorescence and object-level features:

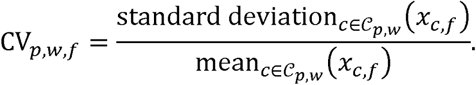

Robust coefficients of variation were calculated for eight cell-shape features using the median absolute deviation:

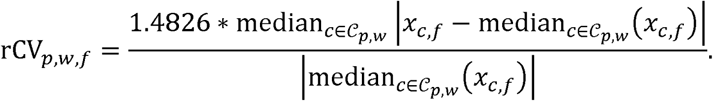

##### Plate- and channel-level quality control

Wild-type (WT) wells, incorporated in each plate, were used to identify large plate-level technical artifacts after profile aggregation and plate-median rescaling. For each selected quality-control feature, the absolute z-score of each WT well relative to the overall WT distribution was calculated as:

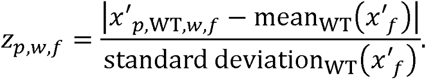

WT wells were flagged when NC ratio, nucleoid-position, nucleoid-fraction, or cell-size-variability features deviated by more than 3 standard deviations from the overall WT mean. A threshold of 5 standard deviations was applied to cell-constriction and cell-volume features. When multiple WT wells on the same plate were flagged for a given feature, the shared deviation was interpreted as a plate-level technical artefact rather than biological variation. The affected feature was therefore set to missing for all samples on that plate (WT and mutant strains). For channel-specific measurements, all features derived from the corresponding fluorescence channel were set to missing when the observed pattern was consistent with a channel-wide staining or imaging artefact. Additional artefacts identified by visual inspection were removed by setting inner membrane features to missing for Δ*fepE*, Δ*mmuM*, Δ*caiC*, Δ*copA*, Δ*caiA*, Δ*metH*, Δ*ydhS*, and Δ*yedJ*, and nucleoid features to missing for Δ*ydeA*.

This preprocessed dataset comprises 4,282 well-level profiles from 94 plates, representing 3,704 unique labels, including WT, with 579 WT wells. The final table contained 699 columns: metadata, 661 original median-aggregated features, 20 CV features, 8 rCV features, and 5 extra heterogeneity features. This dataset (Dataset S3) was used to create Fig. 3 and Extended Data Fig. 3.

##### Normalized score calculations

To detect outliers, we calculated normalized scores (s) for each well-level feature:

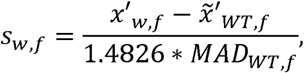

where x’_w,f_ is the measured value of feature *f* in well *w, x^~1^_WT,f_* the median value of that feature across all WT wells, and *MAD_WT,f_* is the corresponding median absolute deviation. This robust normalization preserves whether each feature is increased or decreased relative to the WT while reducing sensitivity to extreme WT values.

##### Normalization and feature selection for profile correlation and embedding analysis

For downstream analyses such as profile correlations and UMAP embeddings, a normalization and feature selection step was performed. Well-level profiles were normalized relative to WT controls using MAD robustization implemented with *pycytominer.normalize*. WT samples from all plates were pooled to estimate the reference median and dispersion for each feature. Each value was centered on the WT median and scaled by the WT median absolute deviation. Non-metadata features were then filtered using *pycytominer.feature_select* to remove outlier features, low-variance features, and highly correlated features. Highly correlated features were filtered using a Pearson correlation threshold of 0.97, with na_cutoff = 0.5, freq_cut = 0.01, unique_cut = 0.1, and outlier_cutoff = 100. This procedure retained 412 features for downstream analysis.

##### Correlation coefficients

Pairwise gene-gene profile similarities were calculated as Spearman rank correlations using the *scipy.stats.spearmanr* method, with missing values omitted. The same procedure was used to calculate feature-feature correlations.

##### Weighted morphological profiling network analysis

An adapted WGCNA framework was used to identify groups of deletion strains with similar morphological profiles. Pairwise Spearman correlations between gene-level profiles were symmetrized and constrained to the interval [-1, 1]; missing or infinite values were set to zero.

#### A signed weighted adjacency matrix was calculated using the soft-thresholding transformation

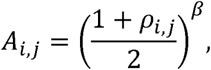

with β = 8. Diagonal entries were set to zero to remove self-edges. Gene connectivity was defined as

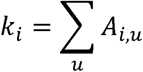

Topological overlap matrix (TOM) was then calculated as

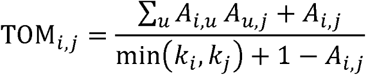

Topological-overlap values were converted to dissimilarities:

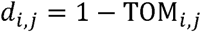

Genes were hierarchically clustered using average linkage, and modules were defined by static tree cutting at *d* = 0.935. Modules containing fewer than three genes were retained as singleton clusters. This procedure yielded 1,545 clusters in total, including 114 multi-gene modules.

##### Annotation of RpoS and ppGpp regulatory associations

Gene-level RpoS and ppGpp annotations were compiled from five studies of *E. coli* K-12. RpoS-evidence comprised transcriptomic and promoter-binding analyses^129,130^, supplemented with expression associations and integrated RNA-seq/ChIP-seq targets^14^. ppGpp-evidence comprised comparisons of ppGpp-deficient cells and controlled ppGpp induction^131,132^. Study-specific selection criteria were retained, and genes with at least one supporting record, including direct or indirect evidence, were classified as associated with RpoS, ppGpp or both. Regulatory effects were labelled activated or repressed when source records agreed, and mixed/condition-dependent when opposing directions were reported. “No documented association” indicated absence of qualifying evidence in the selected studies, rather than demonstrated absence of regulation. Deletion mutants were classified as morphologically divergent when their position in the median profile UMAP space fell outside the contour enclosing 90% of the wild-type probability mass, determined using Gaussian kernel density estimation.

#### Characterization of selected genes of unknown function

##### Data preprocessing and analysis of growth-phase fold-change in cellular fluorescence

The analysis included 19 y-gene fusion strains and a wild-type control, sampled at 12 h (exponential phase) and 24 h (stationary phase), imaged, and processed using BactoScoop. Of the 110,307 cells entering quality control, 108,989 were retained. Filtering excluded morphological outliers with absolute cell length and cell area z-scores ≥ 5 within each sample, followed by removal of fluorescence measurements (C2_cell_mesh_intensity_mean) below the 0.5th or above the 99.5th percentile of the population. After outlier removal, single-cell fluorescence measurements were averaged within each replicate group, and phase-level fluorescence was calculated as the unweighted arithmetic mean of the replicate means. This gave each replicate group equal weight irrespective of its cell count. The log₂ fold change was calculated as:

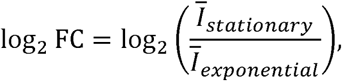

Uncertainty was estimated using 10,000 bootstrap iterations, independently resampling replicate means with replacement within each phase and recalculating the log₂ fold change. The 2.5th and 97.5th percentiles of the bootstrap distribution defined the 95% confidence interval.

Statistical significance was assessed using planned two-sided contrasts from a linear model of log₂-transformed replicate mean fluorescence intensities, including strain, growth phase, and their interaction. Contrasts tested whether the change from exponential to stationary phase differed between each strain and the control. Replicates were treated as independent, and *p*-values were adjusted using Holm’s method, with adjusted *p* < 0.05 considered significant.

##### Retrieval and analysis of published exponential-to-stationary-phase omics data

Gene-level exponential-to-stationary-phase changes in *E. coli* were compiled from seven studies: RNA abundance from Lim et al.^6^, Ekdahl et al.^103^, Morin et al.^104^, Houser et al.^7^ and Lal et al.^105^, protein abundance from Schmidt et al.^8^ and Manzanarez et al.^15^, and mRNA half-lives from Morin et al.^104^ Data were retrieved from supplementary datasets and public repositories and matched by gene name, locus identifier or UniProt accession. Where required, log₂ fold changes were calculated as log₂(stationary/exponential) using means of linear-scale measurements. Published protein fold ratios were log₂-transformed, and SILAC ratios sharing a common reference were divided before transformation. Morin RNA values used linearized stability-assay T0 signals, whereas half-life changes used published fitted estimates. For Manzanarez et al., T1 and T2 denote protein abundance at 6 and 24 h after glucose exhaustion, respectively, each compared with the exponential-phase reference. Positive values indicate increased readout values in stationary phase. Unavailable measurements remained missing without imputation. Genes were ordered by the corresponding fold changes from this study, excluding the wild-type control. Study-specific sources, normalization, endpoints and reliability criteria are detailed in Dataset S8.

##### Conservation analysis of y-genes

Conservation of YeaC, YhcB, YbaB, YoaB, YjgH, YtfB, YggE, YoaC and YebV across the tree of life was assessed using an approach adapted from Sondervorst et al.^52^ Protein sequences were retrieved from the *E. coli* K-12 BW25113 reference genome (CP009273.1) and searched against the archived eggNOG 5.0^133^ proteomes using DIAMOND v2.2.8^134^ in ultra-sensitive mode. Candidate orthologs were evaluated using sequence coverage, reciprocal searches against complete BW25113 reference proteomes, and archived orthology and gene-tree evidence. Supported assignments required an E-value ≤ 10⁻³, ≥ 60% coverage of both proteins, unique reciprocal best matches and additional orthology support. YoaB and YjgH were evaluated separately to distinguish related paralogs. Supported presence was recorded in either the sampled displayed taxon or another sampled species of the same genus; missing sampling and ambiguous assignments remained unresolved. Results were mapped onto an adjusted tree of life (i.e. not containing multiple representatives of the same species) derived from the public iTOL source tree^135^, preserving branch lengths.

The within-species conservation in *E. coli* was done on the gene level. The genes were implemented into a database, which was screened using ABRicate (https://github.com/tseemann/ABRICATE) with a threshold of 90% for query coverage and identity against a large-scale *E. coli* genomic collection of 136,051 genomes^136^.

##### Homology and structural predictions

Homology searches and predictions were performed with the MPI Bioinformatics Toolkit^137^, specifically HMMER^138^ and HHpred^139^. Searches were performed against the PDB_mmCIF70 (HMM-based search and alignment generation against PDB chains; filtered for a maximum sequence identity of 70%) or Pfam-A databases. For more accurate prediction of transmembrane domains and signal peptides, DeepTMHMM 1.0^140^ and SignalP 6.0^141^ were used. The topology diagram was generated using Protter^142^. Structural predictions of y-gene-derived proteins were generated using AlphaFold 3^143^. In cases where y-genes encoded predicted membrane domains (e.g., YhcB), predictions included 40-50 oleic acid (OLA) ligands to stabilize potential lipid-protein interactions^144^. Structural similarity of predicted protein structures with homologous protein structures (as identified by HHpred searches) was quantified using the pairwise structural alignment tool from RCSB PDB^145^.

## Data availability

The data supporting the findings of this study are provided in the Supplementary Datasets. Owing to their large size, the single-cell-level phenotypic profiles and raw microscopy images used to generate Dataset S3 are available from the corresponding author upon reasonable request.

## Code availability

The scripts used to generate the figures in this study are publicly available through the following GitHub repository: https://github.com/Govers-Lab/Steemans_et_al_2026.

## Acknowledgments

We thank Elisa Pierre Despas for providing the msfTq2ox fluorescent protein. We would also like to thank the members of the Govers Lab for the discussions and the critical reading of the manuscript. B.S. was funded by FLOF PhD scholarship of the KU Leuven. K.S. was funded by a doctoral fellowship from the Research Foundation—Flanders (file number 1110925N). M.H. was funded by the European Union under Horizon Europe: MSCA Actions (Project 101209741). K.N. was funded by a junior postdoctoral fellowship from the Research Foundation—Flanders (file number 1251624N). This work was also supported by a start-up grant (STG/21/068) from the KU Leuven Research Fund and by the European Union (ERC Starting Grant BacterialBlueprint 101040800).

## Author contributions

Conceptualization, B.S. and S.K.G.; methodology, B.S. and S.K.G.; software, B.S.; formal analysis, B.S.; investigation, B.S., K.S., M.H., K.N., E.M., and A.M.; data curation, B.S.; writing – original draft, B.S. and S.K.G.; writing – review & editing, B.S., K.S., M.H., and S.K.G.; visualization, B.S. and M.H.; supervision, S.K.G.; project administration, S.K.G.; funding acquisition, S.K.G.

## SUPPLEMENTARY INFORMATION

**Dataset S1** Single-cell measurements and time-course summaries for the OZA001 cells, producing GFP-DnaN, in M9-glucose and LB, including PHATE inputs and coordinates for M9-glucose.

**Dataset S2** Wild-type single-cell profile measurements, PHATE input features, and embedding coordinates.

**Dataset S3** Median-aggregated image-based profiles, normalized scores, and feature definitions for the Keio deletion screen.

**Dataset S4** Spearman correlation matrix and feature labels for the selected features.

**Dataset S5** Gene-profile correlations, UMAP coordinates, network module assignments and annotations, and correlations for previously described stationary-phase actors.

**Dataset S6** Gene annotations for RpoS- and ppGpp-associated regulation, supporting evidence, UMAP coordinates, and gene classifications and counts underlying the regulator-composition analyses.

**Dataset S7** Fluorescence log₂ fold changes between stationary and exponential phases, bootstrap confidence intervals, statistical test results, replicate-group means, and underlying single-cell measurements.

**Dataset S8** Y-gene RNA abundance, protein abundance, fluorescence and mRNA half-life comparisons, including plotted log₂ fold changes, contributing measurements, and source references.

**Dataset S9** Conservation calls for nine y-gene proteins across 161 taxa, including supporting evidence, taxon and protein annotations, and the displayed tree topology and branch lengths.

### SUPPLEMENTARY TEXT 1

To investigate whether genes with an established role in stationary phase also displayed a deviating phenotype in our dataset, we assembled a literature-based set of previously described *E. coli* stationary-phase actors (Dataset S5). The resulting panel captures the major processes repeatedly associated with entry into and survival during stationary phase: the central RpoS response and its regulation (RpoS, Hfq, RssB, IraP, IraD, ClpX, ClpP), global transcriptional and nucleoid regulators (DksA, SspA, Lrp, IhfA, IhfB), DNA protection (Dps), morphological remodeling (BolA), ribosome hibernation and preservation (Rmf, Hpf, RaiA), metabolic regulation, carbon uptake, and storage (ArcA, FadR, SdhA, GlgA, GlgB, GlgC, GlgP, GlgX, Crp, Cra, Icd, PtsG), and stationary-phase maintenance functions including YeaG, Ppk, SurA, SurE, Pcm, Lon, CspB, CspI, and UspA.

In our screen, several previously described stationary-phase-linked genes exhibited deviations in one or more individual morphological features. We first considered the population median cell morphological dimensions (Extended Data Fig. 3a). Deletion of *rpoS*, the central regulator of the stationary-phase response, was associated with an increased cell length, as was deletion of the ribosome-hibernation factor *hpf*. Differences in cell width were observed for a broader set of mutants: *hfq, sspA,* and *lrp* showed a decreased cell width, whereas *cspB*, *crp*, and *clpP* showed an increased cell width. We next examined population-level variation in cell morphological dimensions by comparing the median robust coefficient of variation (rCV) for cell length and width (Extended Data Fig. 3b). Increased rCV cell length was observed for *glgX, glgA,* and *raiA*. Finally, comparison of the population median nucleocytoplasmic (NC) and RNA-to-cytoplasmic (RC) ratios revealed deviations for *lrp, rpoS,* and *clpP* (Extended Data Fig. 3c).

We next asked whether the deletion strains of these previously identified stationary-phase genes showed broader similarities, by comparing their complete image-based profiles. In the UMAP embedding, several strains fell outside the wild-type-rich region, with some occupying shared neighborhoods (Extended Data Fig. 3d). Subsequent comparison of their well-level profiles using Spearman correlation revealed several blocks of positive correlations along the diagonal of the clustered matrix (Extended Data Fig. 3e). These blocks often connected actors with closely related functions. The strongest correlation was observed between Δ*glgP* and Δ*glgX* (ρ = 0.776), with both genes encoding complementary enzymes involved in glycogen degradation. Positive correlations were also observed between the Δ*glgA* and Δ*glgC* (ρ = 0.657; glycogen synthesis) and between Δ*ihfA* and Δ*ihfB* (ρ = 0.609; both subunits of integration host factor). Other positive correlations connected distinct cellular processes through their shared or interdependent regulation. For example, Δ*hfq* and Δ*hpf* showed similar profiles (ρ = 0.640), consistent with the reported contribution of Hfq to *hpf* expression during stationary phase, thereby linking RNA regulation with ribosome hibernation.

---

**Table S1:** Bacterial strains used and constructed in this study.

| Strain | SGL collection number | Reference | Way of construction |
|---|---|---|---|
| *E. coli* OZA001 | 711 | <sup>43</sup> | N/A |
| BW25113 | 479 |  | N/A |
| BW25113 + pKD46 | 1062 | This work | Electroporation of pKD46 into BW25113 |
| *E. coli* K12 MG1655 *ldtA::ldtA_msfTq2ox-FRT_KAN_FRT* | 1099 |  | N/A |
| DH5α pDHL1029 | 665 |  | N/A |
| BW25113 *ybaB::ybaB-msfgfp-frt* | 1125 | This work | Since replacing the stop codon of *ybaB* with the *msfgfp-frt-nptI-frt* cassette would lead to an incomplete start codon for the downstream *recR* gene, the primer tail in this case did contain the last base of *ybaB* to complete the *recR* start codon. Constructed using primers KS146 and KS147, verified using primers KS148 and KS149, excision of *nptI* via pCP20, then sequenced using primers KS148, KS149, KS69, KS70. |
| BW25113 *yeaC::yeaC-msfgfp-frt* | 1123 | This work | Constructed using primers KS158 and KS159, verified using primers KS160 and KS161, excision of *nptI* via pCP20, then sequenced using primers KS160, KS161, KS69, KS70. |
| BW25113 *yebV::yebV-msfgfp-frt* | 1117 | This work | Constructed using primers KS164 and KS165, verified using primers KS166 and KS167, excision of *nptI* via pCP20, then sequenced using primers KS166, KS167, KS69, KS70. |
| BW25113 *yebW::yebW-msfgfp-frt* | 1153 | This work | Constructed using primers KS170 and KS171, verified using primers KS172 and KS173, excision of *nptI* via pCP20, then sequenced using primers KS172, KS173, KS69, KS70. |
| BW25113 *yggE::yggE-msfTq2ox-frt-nptI-frt* | N/A | This work | The *msfTq2ox-frt-nptI-frt* cassette was first amplified from *E. coli* K12 MG1655 *ldtA::ldtA_msfTq2ox-frt-nptI-frt* with short primers (KS218 and KS219) that omit the start codon of *msfTq2ox*, and then amplified using primers KS220 and KS221, verified using primers KS27 and KS28, then sequenced using primers KS7, KS27, KS28, KS250, KS251, KS252. |
| BW25113 *yggL::yggL-msfgfp-frt-nptI-frt* | N/A | This work | Constructed using primers KS236 and KS237, verified using primers KS222 and KS223, then sequenced using primers KS7, KS222, KS223, KS69, KS70, KS250. |
| BW25113 *ynfA::ynfA-msfgfp-frt* | 1135 | This work | Constructed using primers KS182 and KS183, verified using primers KS184 and KS185, excision of *nptI* via pCP20, then sequenced using primers KS184, KS185, KS69, KS70. |
| BW25113 *yoaC::yoaC-msfgfp-frt* | 1119 | This work | Constructed using primers KS194 and KS195, verified using primers KS196 and KS197, excision of *nptI* via pCP20, then sequenced using primers KS196, KS197, KS69, KS70. |
| BW25113 *ysgA::ysgA-msfgfp-frt* | 1145 | This work | Constructed using primers KS206 and KS207, verified using primers KS208 and KS209, excision of *nptI* via pCP20, then sequenced using primers KS208, KS209, KS69, KS70. |
| BW25113 *yqjF::yqjF-msfgfp-frt* | 1220 | This work | Constructed using primers KS198 and KS199, verified using primers KS202 and KS203, excision of *nptI* via pCP20, then sequenced using primers KS202, KS203, KS69, KS70. |
| BW25113 *ybaM::ybaM-msfgfp-frt* | 1211 | This work | Constructed using primers MH308 and MH309. Verified and sequenced using primers MH350 and MH351. |
| BW25113 *ytfB::ytfB-msfgfp-frt* | 1212 | This work | Constructed using primers MH310 and MH311. Verified and sequenced using primers MH352 and MH353. |
| BW25113 *yedK::yedK-msfgfp-frt* | 1213 | This work | Constructed using primers MH312 and MH313. Verified and sequenced using primers MH332 and MH333. |
| BW25113 *yedJ::yedJ-msfgfp-frt* | 1214 | This work | Constructed using primers MH314 and MH315. Verified and sequenced using primers MH334 and MH335. |
| BW25113 *yoaB::yoaB-msfgfp-frt* | 1215 | This work | Constructed using primers MH316 and MH317. Verified and sequenced using primers MH336 and MH337. |
| BW25113 *yhcB::yhcB-msfgfp-frt* | 1216 | This work | Constructed using primers MH318 and MH319. Verified and sequenced using primers MH338 and MH339. |
| BW25113 *ychF::ychF-msfgfp-frt* | 1217 | This work | Constructed using primers MH320 and MH321. Verified and sequenced using primers MH354 and MH355. |
| BW25113 *yidF::yidF-msfgfp-frt* | 1218 | This work | Constructed using primers MH322 and MH323. Verified and sequenced using primers MH342 and MH343. |
| BW25113 *yjgH::yjgH-msfgfp-frt* | 1219 | This work | Constructed using primers MH326 and MH327. Verified and sequenced using primers MH346 and MH347. |

**Table S2:**
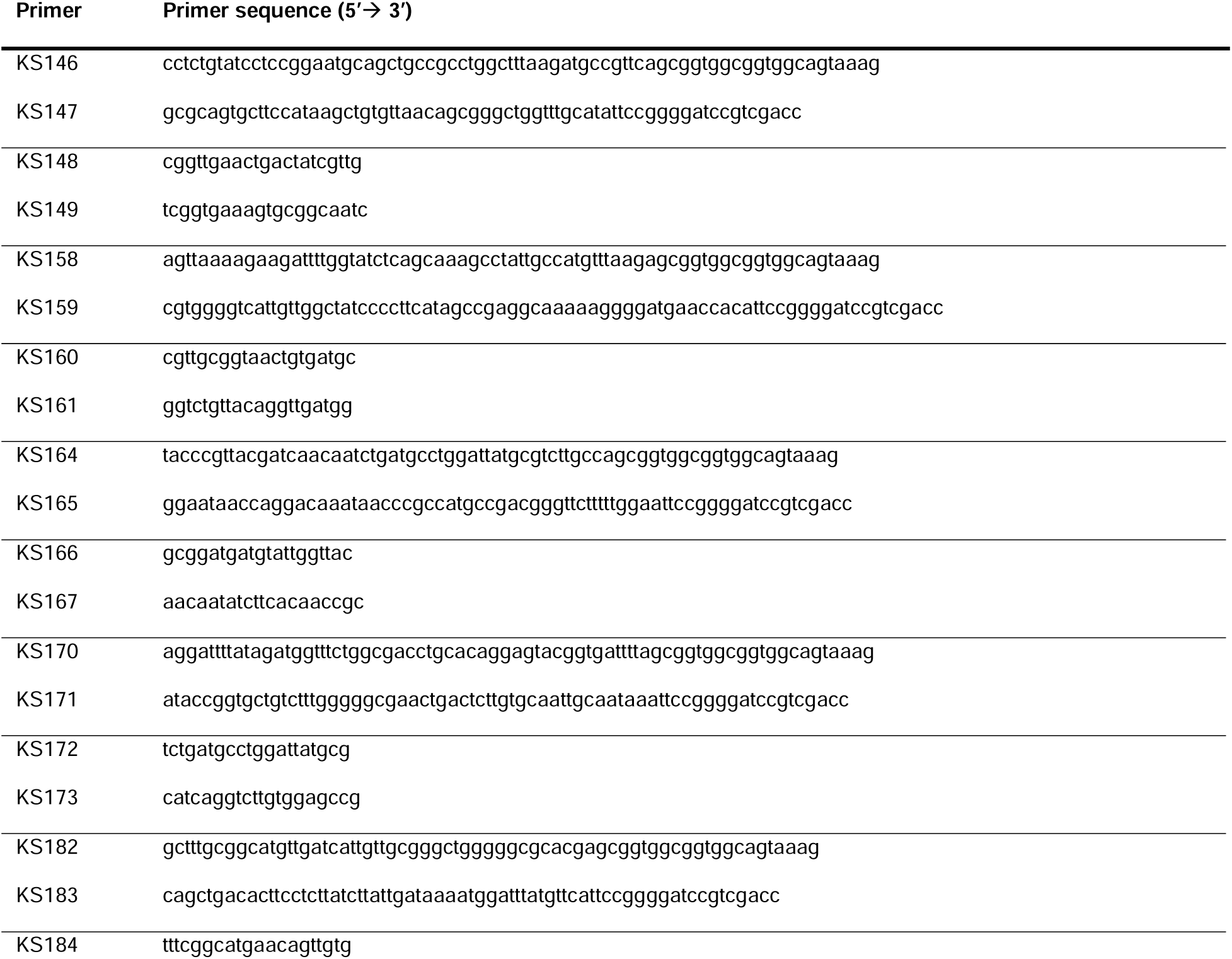
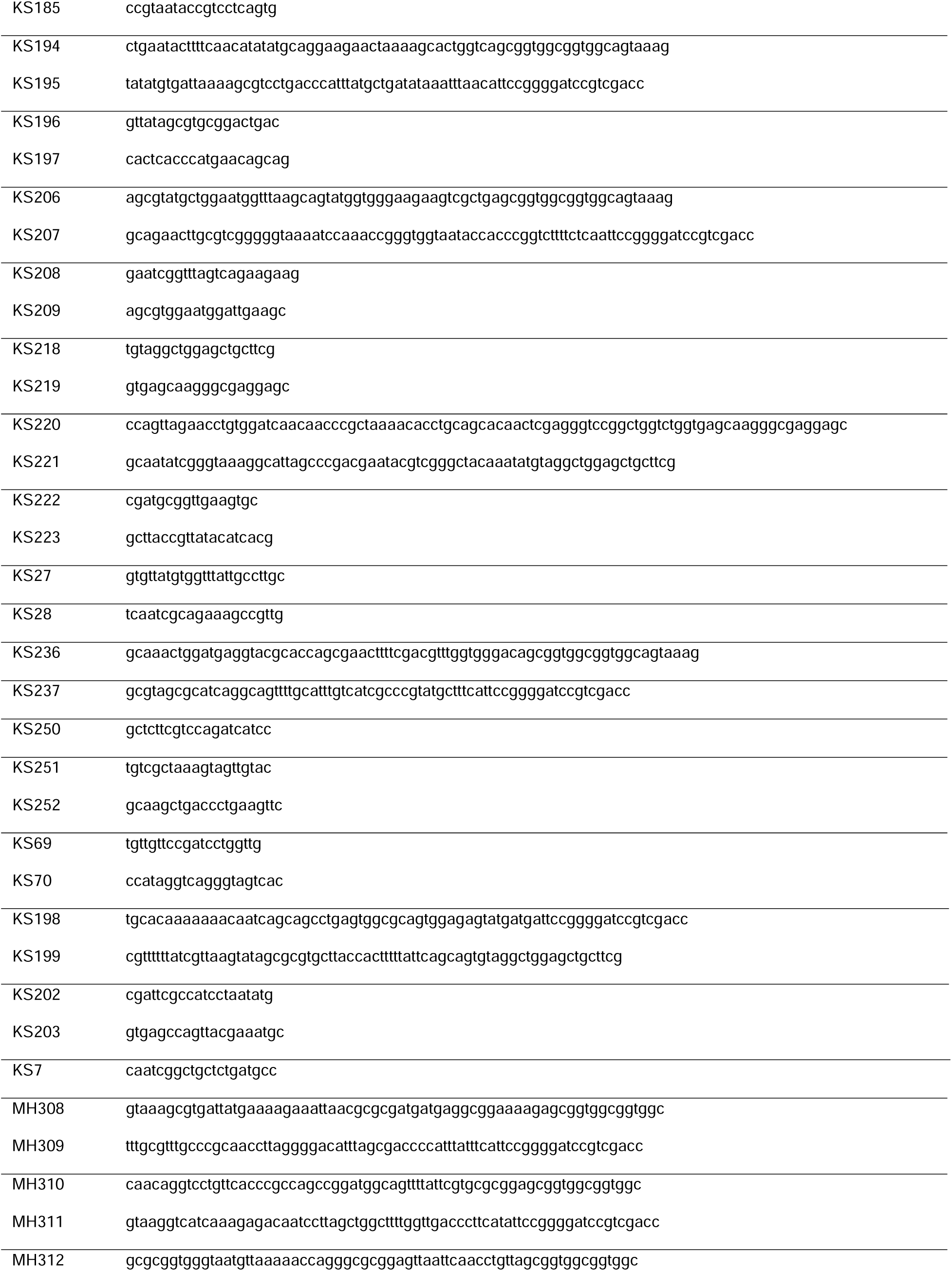
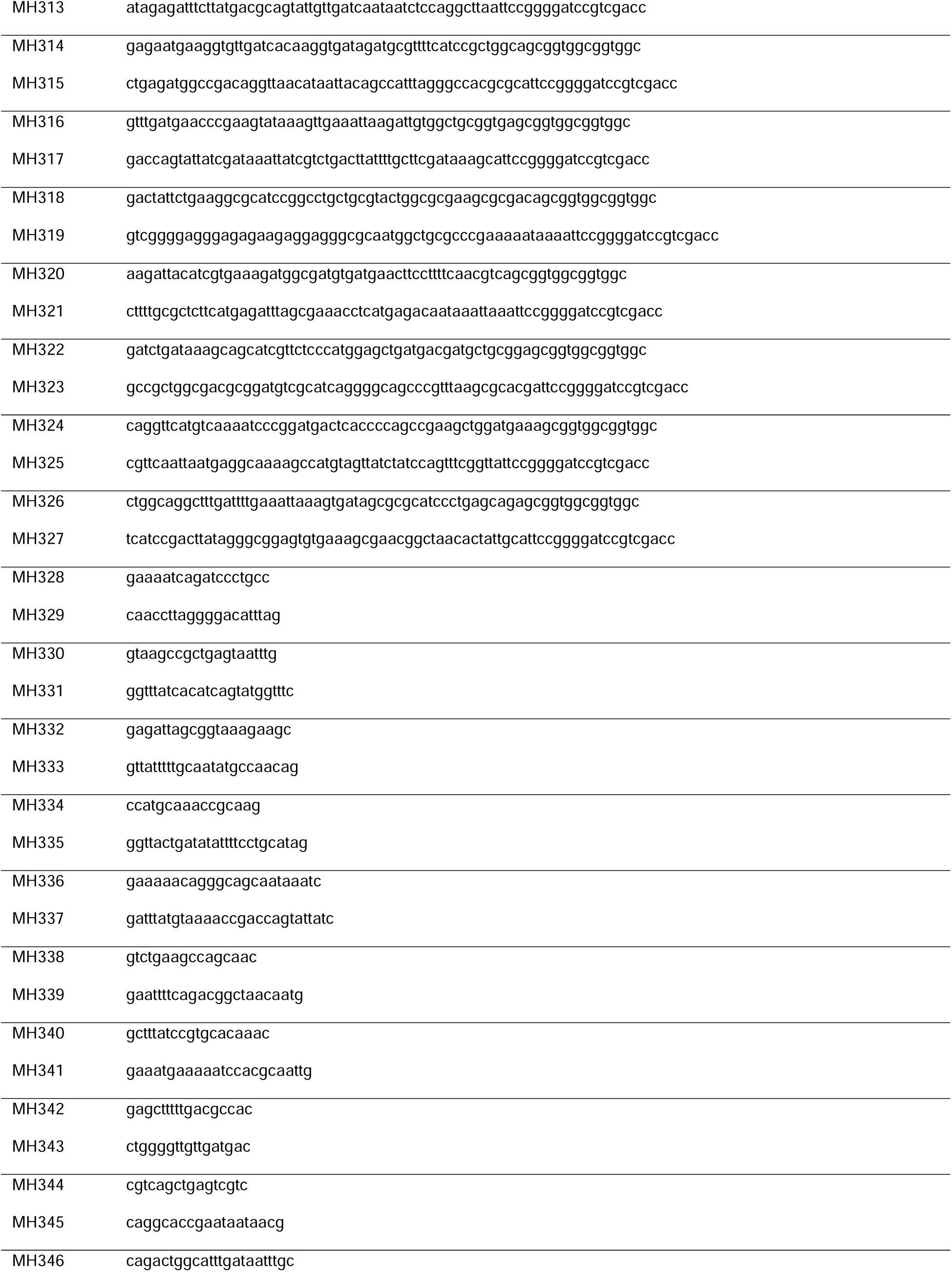
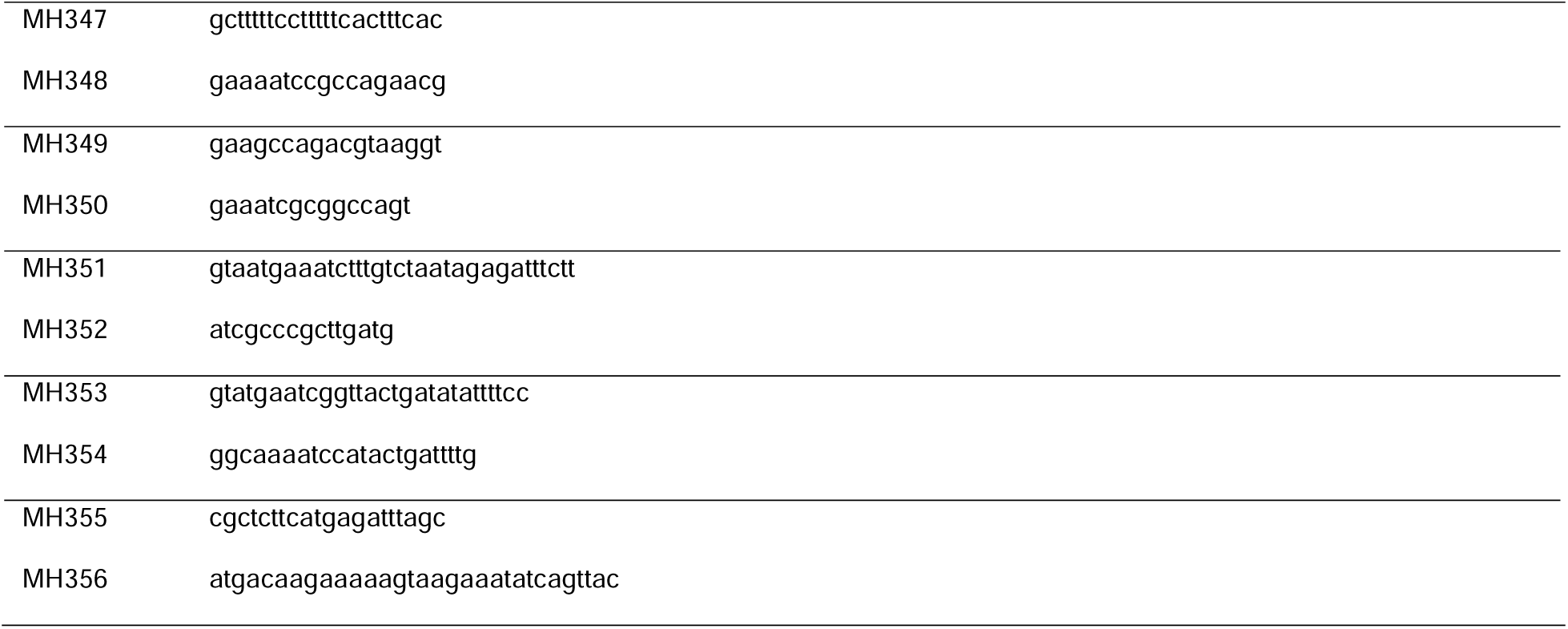
List of primers used in this study.

## EXTENDED DATA FIGURES

**Extended Data Fig. 1: Quantification of cell constriction and chromosome replication, and morphological changes during growth in M9 glucose and LB media. a**, Scatter plot showing the constriction degree (calculated by multiplying the maximal inverted phase-contrast indentation by the maximal width profile indentation) versus the cell length. The data are from *E. coli* OZA001 cells sampled after 12 hours of growth in M9glu, with representative phase-contrast images of a constricting cell (orange) and non-constricting cell (blue). The brown line marks the empirical indentation threshold used to determine the fraction of constricting cells (0.2 after rescaling, equivalent to 0.1 on the original scale). **b**, Scatter plot showing the relative GFP-DnaN projected signal area versus cell volume, with representative GFP-DnaN signal images of a cell performing chromosome replication (blue) and a cell without active chromosome replication (orange). A histogram of the relative frequency of the relative GFP-DnaN area is shown in grey. The brown line at 0.4 indicates the threshold chosen to determine the fraction of cells performing chromosome replication. **c**, Plots showing the behavior of average cell morphological features (cell width, area, surface area, and surface-area-to-volume ratio) over time. Error bars indicate standard deviation between replicates (n ≥ 3). **d**, Representative phase-contrast and epifluorescence images of *E. coli* cells expressing the GFP-DnaN protein grown in LB medium at 3, 6, and 30 h (see Methods). **e,** Changes in mean cell morphological features (length, volume, and aspect ratio) and replicative characteristics (fraction of constricting cells, cells containing a single nucleoid, and cells undergoing active chromosome replication) over time in LB-grown *E. coli* GFP-DnaN populations. Error bars represent the standard deviation across biological replicates (n ≥ 3).

**Extended Data Fig. 2: Reproducibility of stationary-phase single-cell profiles across wild-type replicates.** Twelve scatterplots of the PHATE embedding shown in Fig. 2c, each highlighting one of the 12 wild-type biological replicates against the remaining replicates. The consistent distribution of cells across the embedding illustrates the reproducibility of stationary-phase phenotypic profiles between replicates.

**Extended Data Fig. 3: Image-based phenotypes and profile similarities of previously identified stationary-phase actors. a,** Median cell width versus median cell length, as in Fig. 3a. **b,** Robust coefficient of variation (rCV) in cell width versus rCV in cell length, as in Fig. 3b. **c,** Median RNA-to-cytoplasmic (RC) ratio versus median nucleocytoplasmic (NC) ratio, as in Fig. 3d. In a–c, deviating deletion strains (|s| ≥ 4) of genes that were previously implicated in stationary-phase physiology are highlighted, and black density contours enclose 10%, 30%, 50%, 70%, and 90% of the WT distribution. **d,** UMAP embedding of mutant profiles based on 412 selected features, as in Fig. 5b, with deviating deletion strains highlighted. Black density contours enclose 10%, 25%, 50%, 75%, and 90% of the wild-type distribution. **e,** Heatmap showing pairwise Spearman correlation coefficients (ρ) between the feature profiles of the 38 deletion strains of genes that were previously implicated in stationary-phase physiology. Genes are ordered by hierarchical clustering using the distance 1 − ρ.

**Extended Data Fig. 4: Variation in nucleocytoplasmic and RNA-cytoplasmic ratios is primarily associated with nucleoid and RNA area (and not cell area). a**, Scatter plots showing the nucleocytoplasmic (NC) ratio versus cell area (left) and nucleoid area (right) across wild-type and deletion strains. **b**, Scatterplots showing the RNA-cytoplasmic (RC) ratio versus cell area (left) and RNA area (right) across wild-type and deletion strains. Black density contours indicate regions enclosing 10%, 30%, 50%, 70%, and 90% of the wild-type distribution. Spearman correlation coefficients (ρ) are indicated in each panel.

**Extended Data Fig. 5: Distribution of morphological and subcellular features across the genome-wide stationary-phase phenotypic landscape.** UMAP embedding from Fig. 5b, colored according to six selected features: cell length, cell width, nucleocytoplasmic (NC) ratio, mean RNA fluorescence intensity, mean inner-membrane (IM) fluorescence intensity, and outer-membrane (OM) signal uniformity. Each panel displays the same embedding, highlighting how individual morphological and subcellular features vary across the phenotypic landscape.

**Extended Data Fig. 6: Phenotypic signatures of RNA-turnover-related gene deletion strains Δ*rppH* and Δ*pnp*. a**, Representative phase-contrast and fluorescence images of wild-type (WT), Δ*rppH*, and Δ*pnp* cells, showing the nucleoid (DAPI), RNA (RNASelect), inner membrane staining (MitoTracker Red CMXRos), and outer membrane staining (FM4-64). **b**, Histograms comparing the distributions of cell length, cell width, nucleocytoplasmic ratio, and RNA-cytoplasmic ratio between Δ*rppH* cells and wild-type cells from the same experimental plate. **c**, Corresponding distributions for Δ*pnp* cells compared with wild-type cells from the same plate. The mean (μ) and standard deviation (σ) of each distribution are indicated in the plots in panels **b** and **c**.

**Extended Data Fig. 7: Stationary-phase regulatory associations across morphological phenotypes. a**, Regulatory composition of all mutant profiles (*n* = 3,703), profiles outside the 90% wild-type (WT) probability density contour (*n* = 1,550) and profiles inside the contour (*n* = 2,153). Categories indicate documented associations with RpoS only, ppGpp only, both regulators or neither regulator in the selected evidence collection. **b**, UMAP embeddings colored by stationary-phase regulatory association, RpoS effect and ppGpp effect. Each point represents a median profile, with WT controls shown in black. The contour encloses 90% of the estimated WT probability mass.

**Extended Data Fig. 8: Representative microscopy images of fluorescent protein fusions of poorly characterized *E. coli* genes.** Representative phase-contrast and epifluorescence images of *E. coli* BW25113 and the following fluorescent fusion strains, arranged in ascending order of their log₂ fold change in mean fluorescence intensity between exponential and stationary phase: *yeaC-msfgfp*, *ysgA-msfgfp*, *ychF-msfgfp*, *yqjF-msfgfp*, *ynfA-msfgfp*, *yidF-msfgfp*, *ybaM-msfgfp*, *yedK-msfgfp*, *yggL-msfgfp*, *yedJ-msfgfp*, *yebW-msfgfp*, *yhcB-msfgfp*, *ybaB-msfgfp*, *yoaB-msfgfp*, *ytfB-msfgfp*, *yjgH-msfgfp*, *yggE-msfTq2ox*, *yoaC-msfgfp*, and *yebV-msfgfp*.

**Extended Data Fig. 9: Conservation of the investigated genes within *E. coli* and across the species-labelled tree of life. a**, Bar plot indicating the prevalence of the selected y-genes across a collection of 136,051 *E. coli* genomes. **b**, Species-labelled version of Fig. 6c, showing the distribution of orthologs of the indicated proteins across the tree of life. Symbols are as described in Fig. 6c.

**Extended Data Fig. 10: Structural characterization of selected *E. coli* proteins and associated complexes. a**, AlphaFold 3-predicted structures of YhcB, YbaB, YoaB, YtfB, YjgH, YggE, YoaC, YebV, and YeaC. **b**, Left, AlphaFold 3-predicted structure of the large ACC protein complex (PDB: 9E4O) comprising AccA, AccB (BCCP), AccC, and AccD. Right, cross-section of the AlphaFold3-predicted ACC complex with the predicted YhcB-AccA interaction. **c**, Comparison of an AlphaFold 3-predicted structure of the YggE-complex with an experimentally determined crystal structure of the BP26 protein from *Brucella abortus* (PDB: 4HVZ), obtained by X-ray crystallography at 3.5 Å resolution. Right, colored by MLP. **d**, Top, cross-sectional view of the predicted YggE octamer showing the dimensions of the internal channel. Bottom, top view of the complex showing the central pore.

