## Supplementary material for "Mapping the genetic landscape of cell morphogenesis and subcellular organization in stationary-phase *Escherichia coli* cells": Figure S1

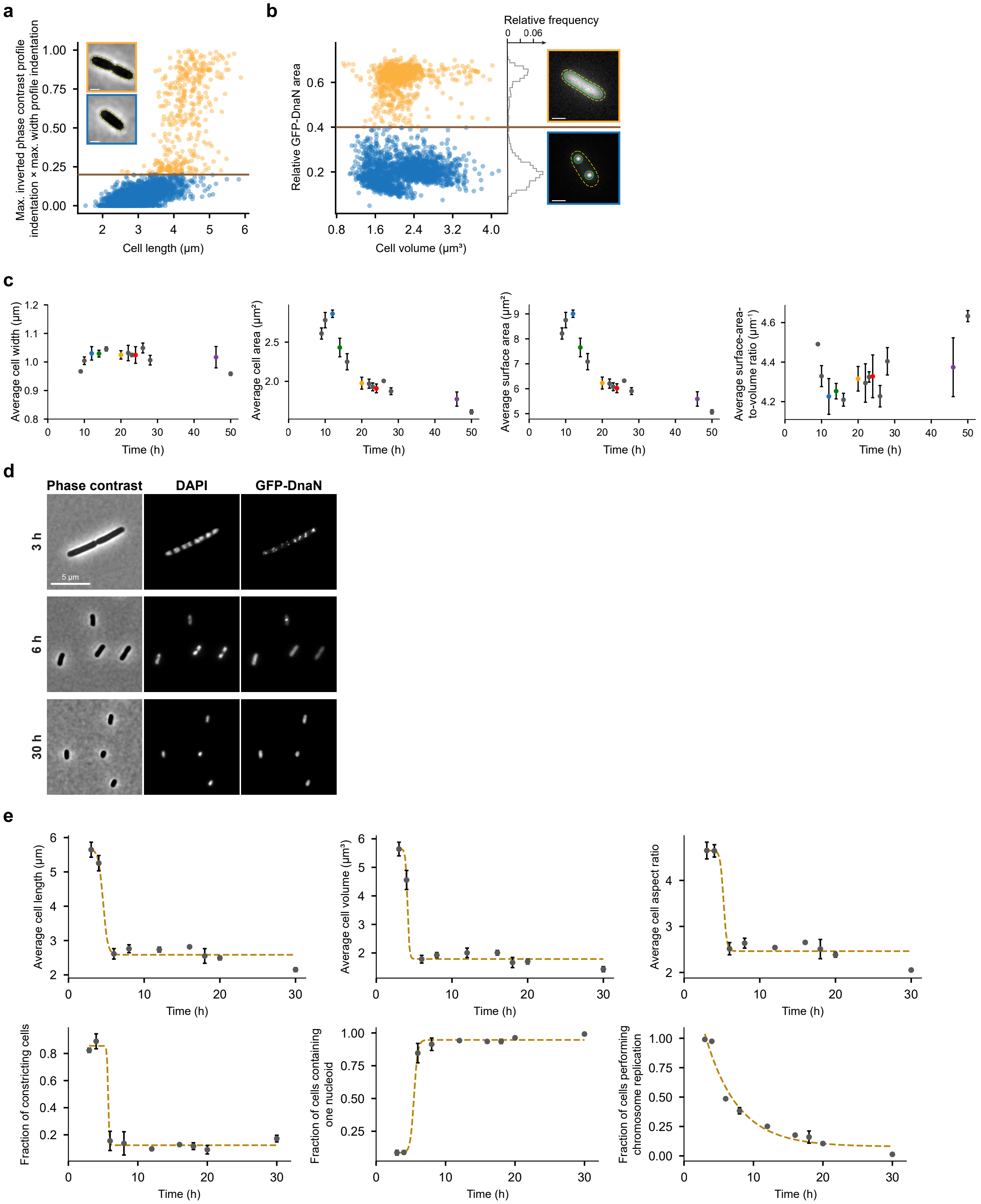

a
Max. inverted phase contrast profile indentation × max. width profile indentation
Cell length (μm)
b
Relative frequency
0
0.06
Relative GFP-DnaN area
Cell volume (μm³)
c
Average cell width (μm)
Average cell area (μm²)
Average surface area (μm²)
Average surface-area-to-volume ratio (μm⁻¹)
Time (h)
d
Phase contrast
DAPI
GFP-DnaN
3 h
6 h
30 h
5 μm
e
Average cell length (μm)
Average cell volume (μm³)
Average cell aspect ratio
Fraction of constricting cells
Fraction of cells containing one nucleoid
Fraction of cells performing chromosome replication
Time (h)
