## Supplementary material for "Mapping the genetic landscape of cell morphogenesis and subcellular organization in stationary-phase *Escherichia coli* cells": Figure S2

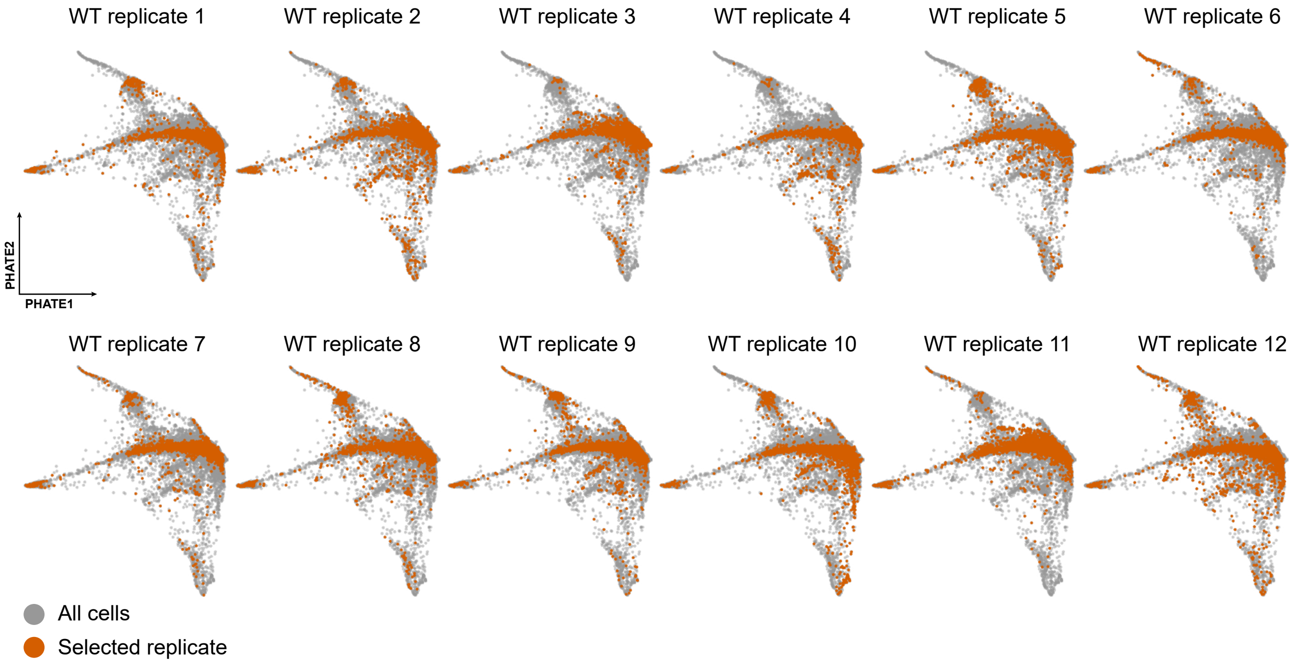
WT replicate 1
WT replicate 2
WT replicate 3
WT replicate 4
WT replicate 5
WT replicate 6
PHATE2
PHATE1
WT replicate 7
WT replicate 8
WT replicate 9
WT replicate 10
WT replicate 11
WT replicate 12
All cells
Selected replicate
