## Supplementary material for "Mapping the genetic landscape of cell morphogenesis and subcellular organization in stationary-phase *Escherichia coli* cells": Figure S3

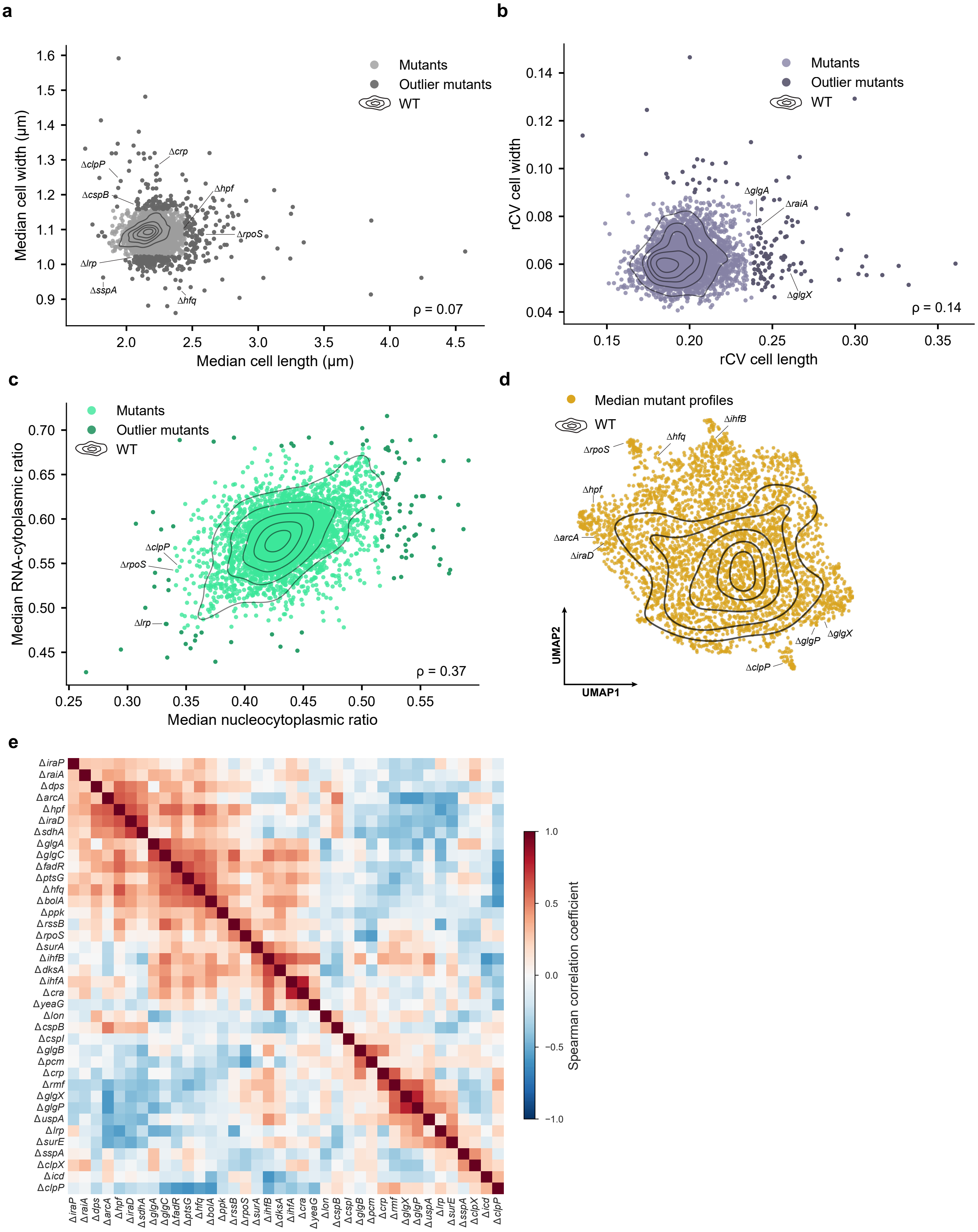
a
Mutants
Outlier mutants
WT
Median cell width (µm)
Median cell length (µm)
ΔclpP
ΔcspB
Δcrp
Δhpf
ΔrpoS
Δlrp
ΔsspA
Δhfq
ρ = 0.07
b
Mutants
Outlier mutants
WT
rCV cell width
rCV cell length
ΔglgA
ΔraiA
ΔglgX
ρ = 0.14
c
Mutants
Outlier mutants
WT
Median RNA-cytoplasmic ratio
Median nucleocytoplasmic ratio
ΔclpP
ΔrpoS
Δlrp
ρ = 0.37
d
Median mutant profiles
WT
ΔihfB
Δhfq
ΔrpoS
Δhpf
ΔarcA
ΔiraD
ΔglgX
ΔglgP
ΔclpP
UMAP2
UMAP1
e
ΔiraP
ΔraiA
Δdps
ΔarcA
Δhpf
ΔiraD
ΔsdhA
ΔglgA
ΔglgC
ΔfadR
ΔptsG
Δhfq
ΔbolA
Δppk
ΔrssB
ΔrpoS
ΔsurA
ΔihfB
ΔdksA
ΔihfA
Δcra
ΔyeaG
Δlon
ΔcspB
ΔcspI
ΔglgB
Δpcm
Δcrp
Δrmf
ΔglgX
ΔglgP
ΔuspA
Δlrp
ΔsurE
ΔsspA
ΔclpX
Δicd
ΔclpP
Spearman correlation coefficient
1.0
0.5
0.0
−0.5
−1.0
