## Supplementary material for "Mapping the genetic landscape of cell morphogenesis and subcellular organization in stationary-phase *Escherichia coli* cells": Figure S4

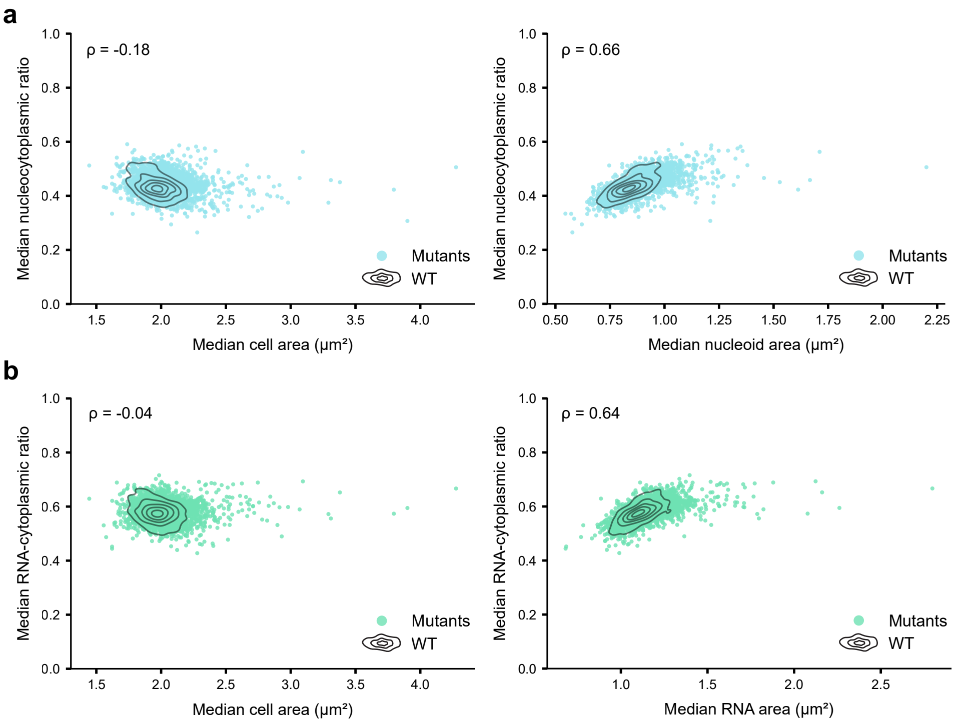
a
ρ = -0.18
Median nucleocytoplasmic ratio
Median cell area (µm²)
Mutants
WT
ρ = 0.66
Median nucleocytoplasmic ratio
Median nucleoid area (µm²)
Mutants
WT
b
ρ = -0.04
Median RNA-cytoplasmic ratio
Median cell area (µm²)
Mutants
WT
ρ = 0.64
Median RNA-cytoplasmic ratio
Median RNA area (µm²)
Mutants
WT
