## Supplementary material for "Mapping the genetic landscape of cell morphogenesis and subcellular organization in stationary-phase *Escherichia coli* cells": Figure S5

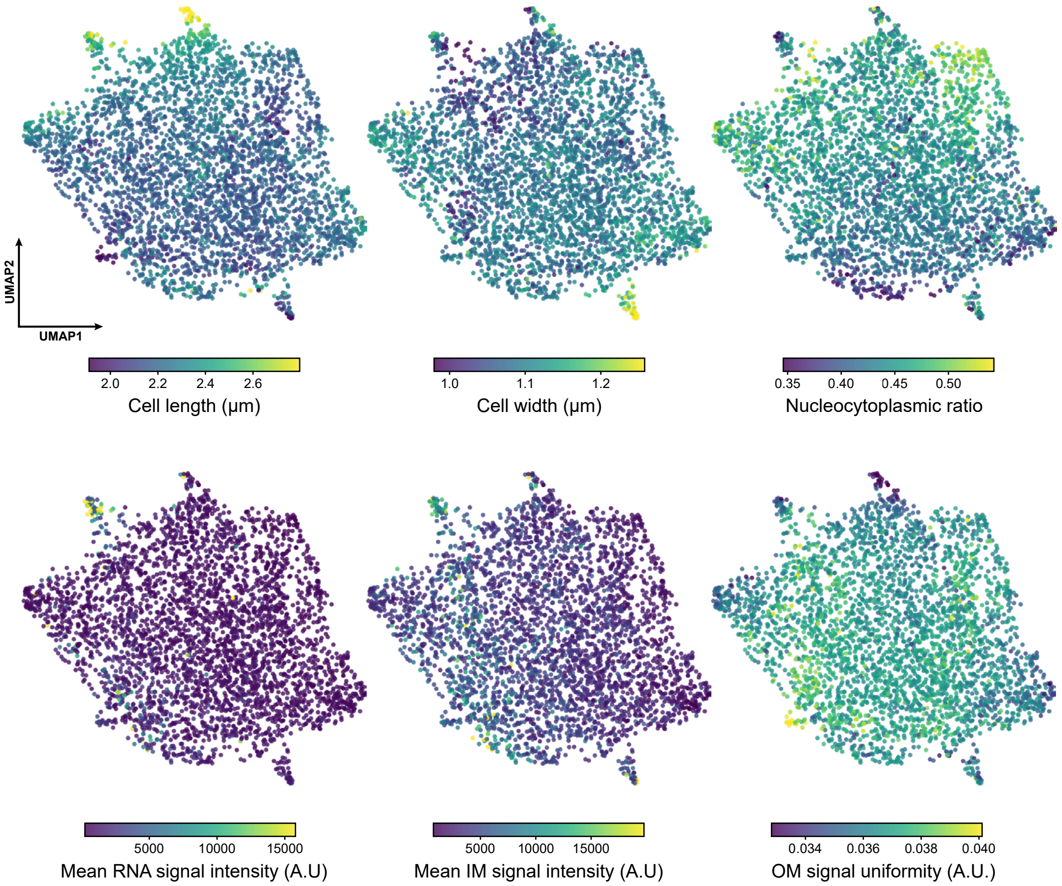
UMAP2
UMAP1
2.0 2.2 2.4 2.6
Cell length (µm)
1.0 1.1 1.2
Cell width (µm)
0.35 0.40 0.45 0.50
Nucleocytoplasmic ratio
5000 10000 15000
Mean RNA signal intensity (A.U)
5000 10000 15000
Mean IM signal intensity (A.U)
0.034 0.036 0.038 0.040
OM signal uniformity (A.U.)
