## Supplementary material for "Mapping the genetic landscape of cell morphogenesis and subcellular organization in stationary-phase *Escherichia coli* cells": Figure S6

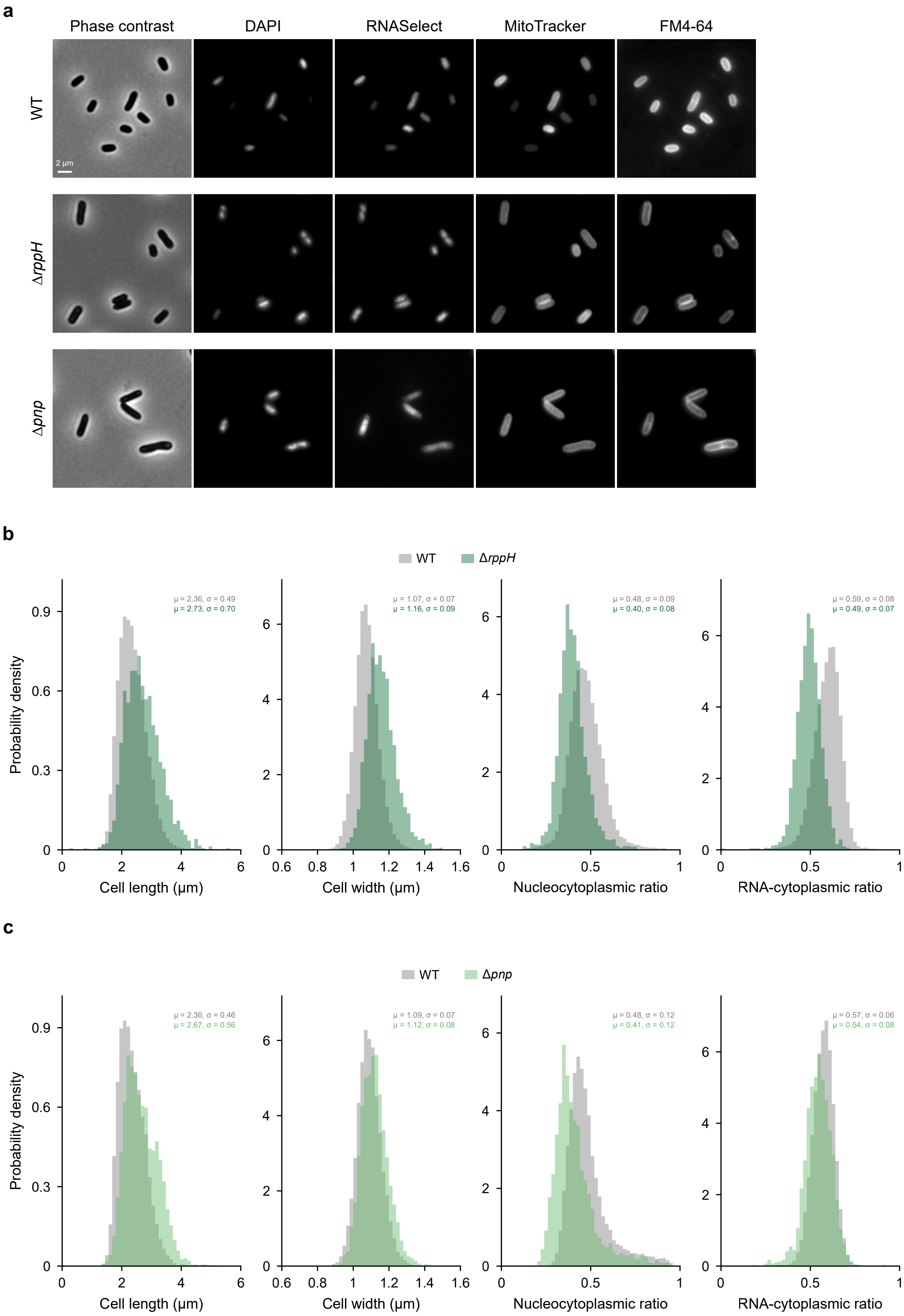
a
Phase contrast
DAPI
RNASelect
MitoTracker
FM4-64
WT
ΔrppH
Δpnp
2 μm
b
WT
ΔrppH
μ = 2.36, σ = 0.49
μ = 2.73, σ = 0.70
μ = 1.07, σ = 0.07
μ = 1.16, σ = 0.09
μ = 0.48, σ = 0.09
μ = 0.40, σ = 0.08
μ = 0.59, σ = 0.08
μ = 0.49, σ = 0.07
Probability density
Cell length (μm)
Cell width (μm)
Nucleocytoplasmic ratio
RNA-cytoplasmic ratio
c
WT
Δpnp
μ = 2.36, σ = 0.46
μ = 2.67, σ = 0.56
μ = 1.09, σ = 0.07
μ = 1.12, σ = 0.08
μ = 0.48, σ = 0.12
μ = 0.41, σ = 0.12
μ = 0.57, σ = 0.06
μ = 0.54, σ = 0.08
Probability density
Cell length (μm)
Cell width (μm)
Nucleocytoplasmic ratio
RNA-cytoplasmic ratio
