## Supplementary material for "Mapping the genetic landscape of cell morphogenesis and subcellular organization in stationary-phase *Escherichia coli* cells": Figure S7

**a**

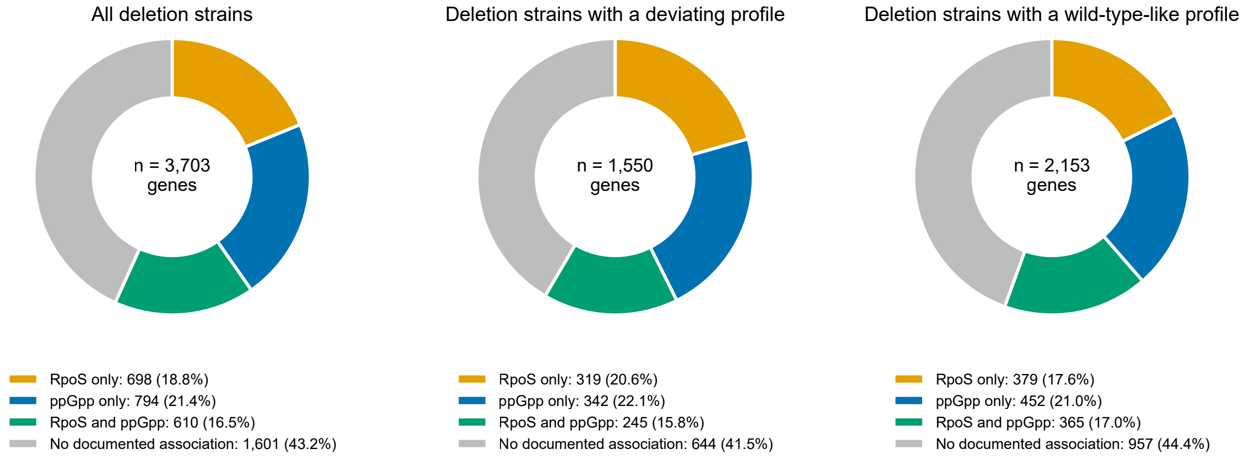
All deletion strains
n = 3,703 genes
Deletion strains with a deviating profile
n = 1,550 genes
Deletion strains with a wild-type-like profile
n = 2,153 genes
RpoS only: 698 (18.8%)
ppGpp only: 794 (21.4%)
RpoS and ppGpp: 610 (16.5%)
No documented association: 1,601 (43.2%)
RpoS only: 319 (20.6%)
ppGpp only: 342 (22.1%)
RpoS and ppGpp: 245 (15.8%)
No documented association: 644 (41.5%)
RpoS only: 379 (17.6%)
ppGpp only: 452 (21.0%)
RpoS and ppGpp: 365 (17.0%)
No documented association: 957 (44.4%)

**b**

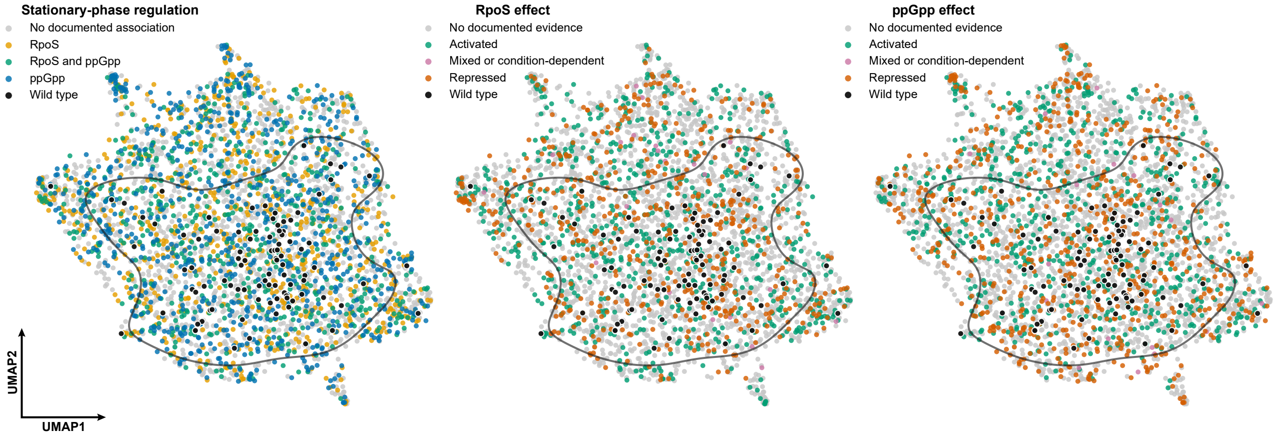
Stationary-phase regulation
No documented association
RpoS
RpoS and ppGpp
ppGpp
Wild type
RpoS effect
No documented evidence
Activated
Mixed or condition-dependent
Repressed
Wild type
ppGpp effect
No documented evidence
Activated
Mixed or condition-dependent
Repressed
Wild type
UMAP2
UMAP1
