## Supplementary material for "Mapping the genetic landscape of cell morphogenesis and subcellular organization in stationary-phase *Escherichia coli* cells": Figure S8

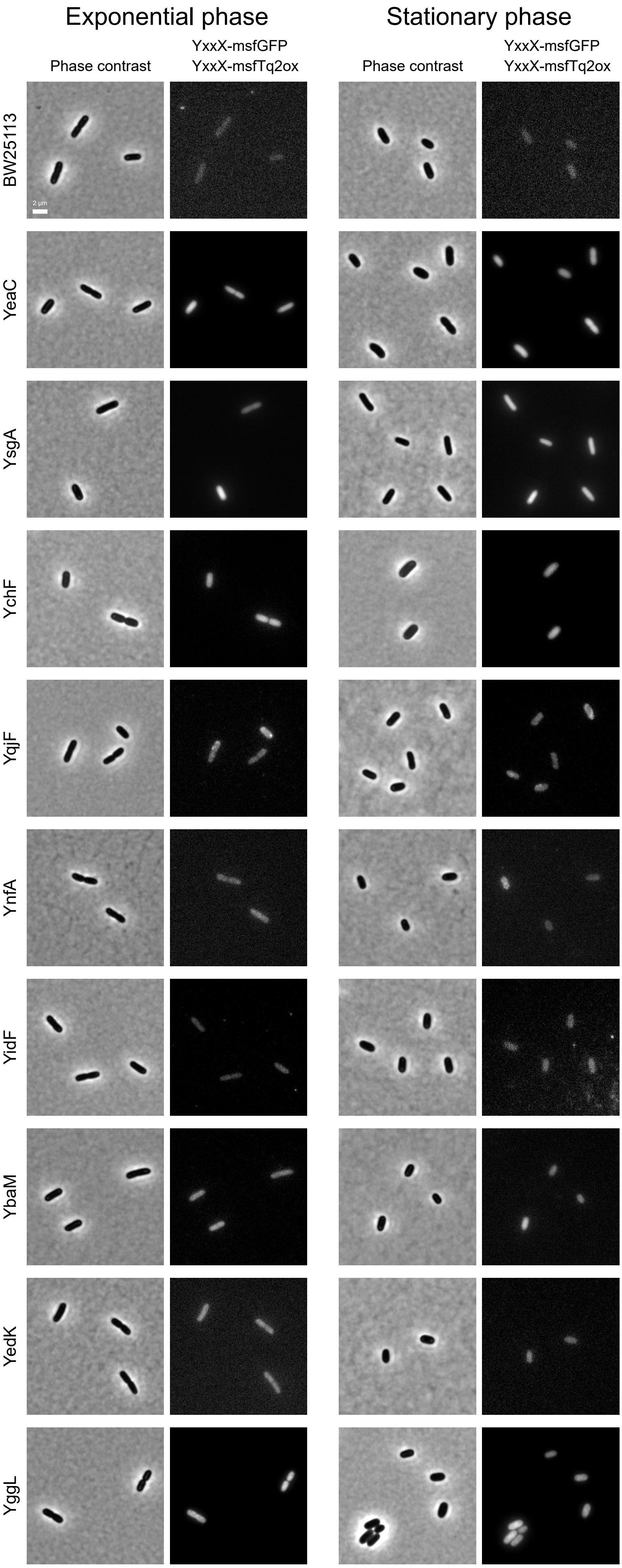

Exponential phase
Stationary phase
Phase contrast
YxxX-msfGFP
YxxX-msfTq2ox
Phase contrast
YxxX-msfGFP
YxxX-msfTq2ox
BW25113
2 µm
YeaC
YsgA
YchF
YqjF
YnfA
YidF
YbaM
YedK
YggL

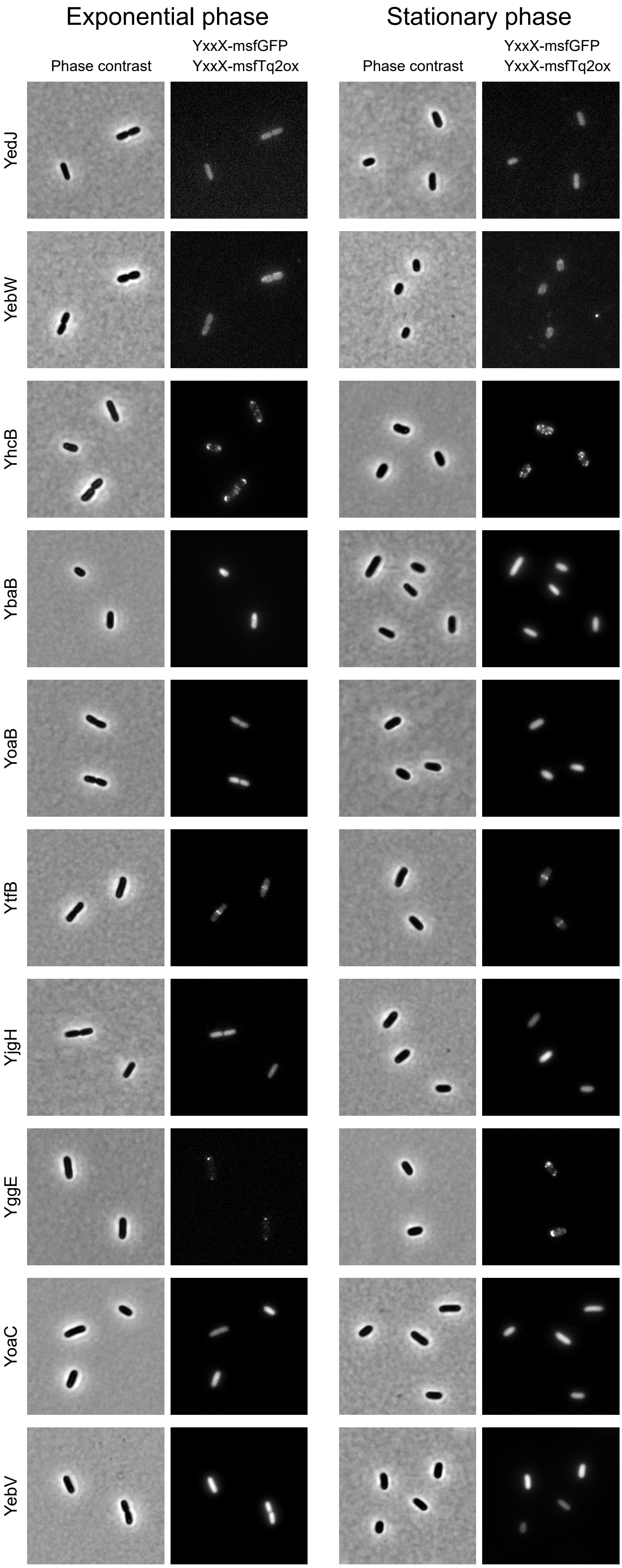

Exponential phase
Stationary phase
Phase contrast
YxxX-msfGFP
YxxX-msfTq2ox
Phase contrast
YxxX-msfGFP
YxxX-msfTq2ox
YedJ
YebW
YhcB
YbaB
YoaB
YtfB
YjgH
YggE
YoaC
YebV
