## Supplementary material for "Mapping the genetic landscape of cell morphogenesis and subcellular organization in stationary-phase *Escherichia coli* cells": Figure S9

**a**

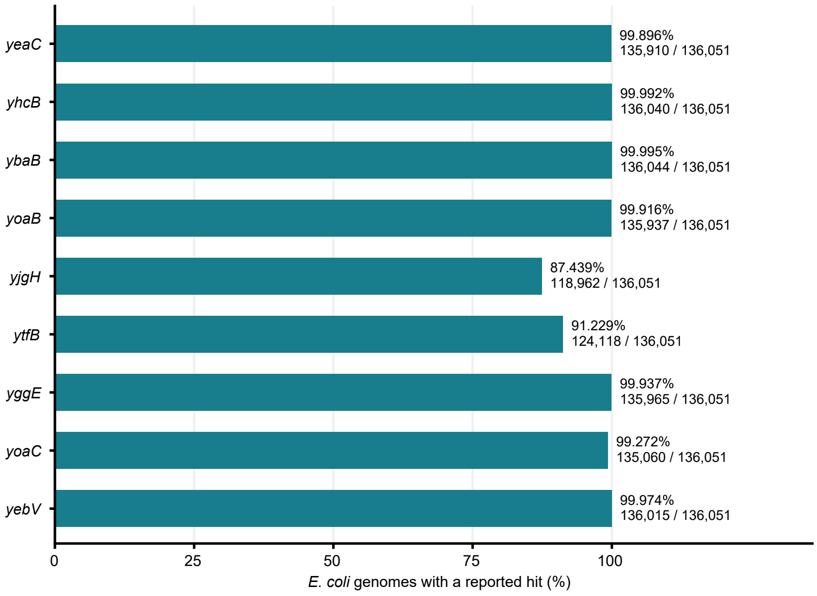
yeaC
yhcB
ybaB
yoaB
yjgH
ytfB
yggE
yoaC
yebV
99.896%
135,910 / 136,051
99.992%
136,040 / 136,051
99.995%
136,044 / 136,051
99.916%
135,937 / 136,051
87.439%
118,962 / 136,051
91.229%
124,118 / 136,051
99.937%
135,965 / 136,051
99.272%
135,060 / 136,051
99.974%
136,015 / 136,051
0
25
50
75
100
E. coli genomes with a reported hit (%)

**b**

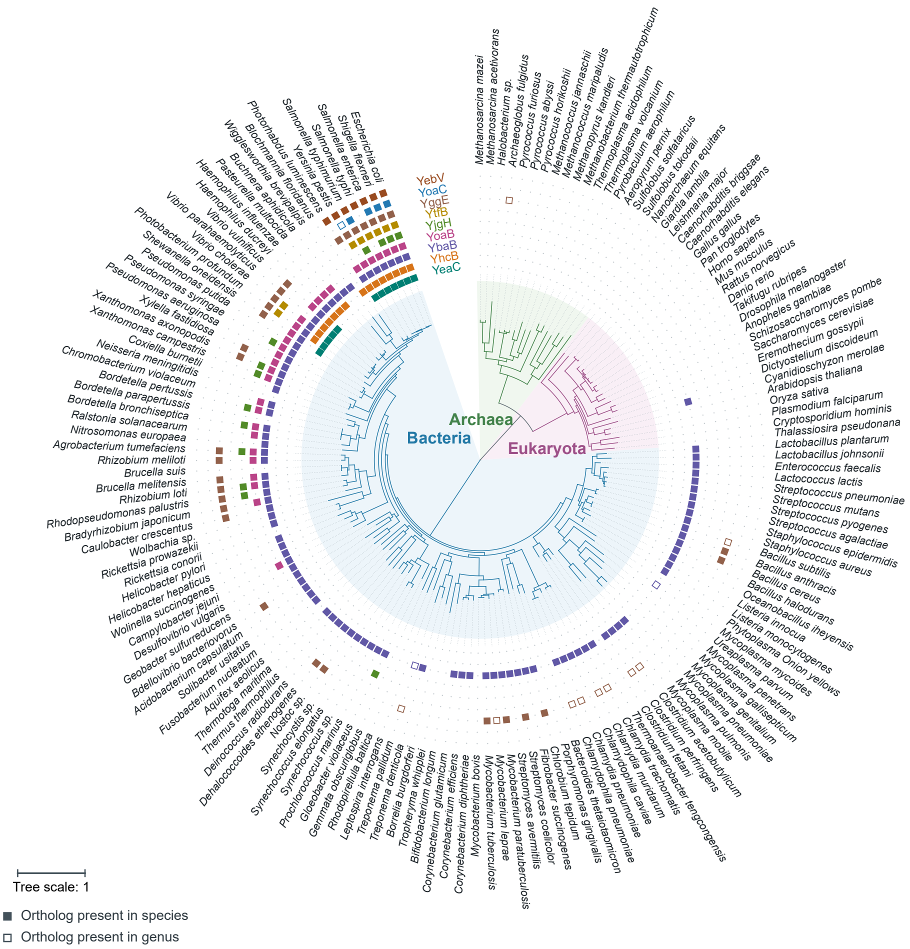
YebV
YoaC
YggE
YtfB
YjgH
YoaB
YbaB
YhcB
YeaC
Archaea
Bacteria
Eukaryota
Tree scale: 1
Ortholog present in species
Ortholog present in genus
