## Supplementary material for "Mapping the genetic landscape of cell morphogenesis and subcellular organization in stationary-phase *Escherichia coli* cells": Figure S10

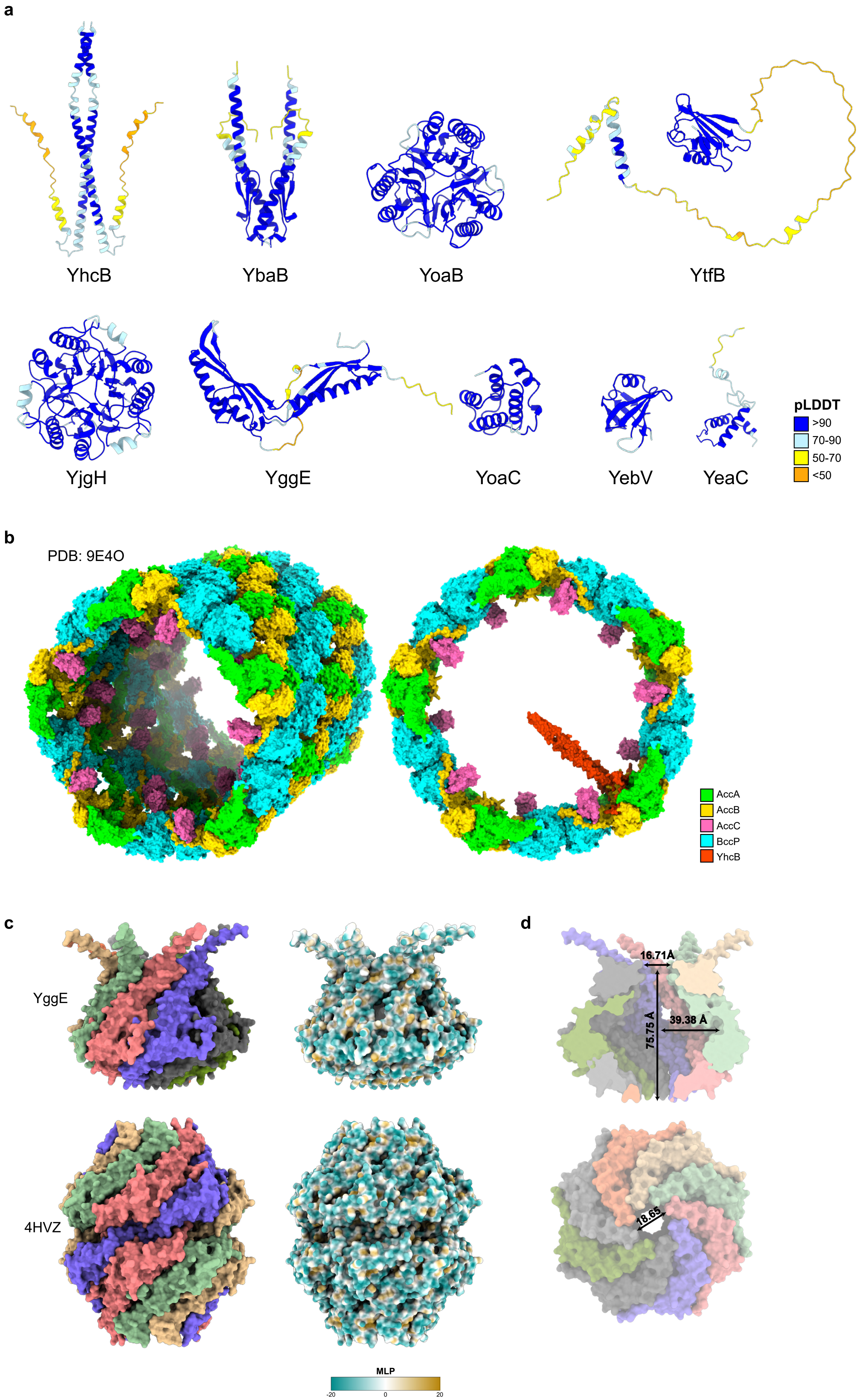

a
YhcB
YbaB
YoaB
YtfB
YjgH
YggE
YoaC
YebV
YeaC
pLDDT
>90
70-90
50-70
<50
b
PDB: 9E4O
AccA
AccB
AccC
BccP
YhcB
c
YggE
4HVZ
MLP
-20
0
20
d
16.71Å
75.75 Å
39.38 Å
18.65
